# Maturational trajectories of white matter microstructure underlying the right presupplementary motor area reflect individual improvements in motor response cancellation in children and adolescents

**DOI:** 10.1101/2020.03.13.990507

**Authors:** Kathrine Skak Madsen, Louise Baruël Johansen, Wesley K. Thompson, Hartwig R. Siebner, Terry L. Jernigan, William F. C. Baaré

**Author notes:** Correspondence: Kathrine Skak Madsen, Danish Research Centre for Magnetic Resonance, Centre for Functional and Diagnostic Imaging and Research, Copenhagen University Hospital Hvidovre, Kettegaard Alle 30, 2650 Hvidovre, Denmark.

## Abstract

The ability to effectively suppress motor response tendencies is essential for focused and goal-directed behavior. Here, we tested the hypothesis that developmental improvement in the ability to cancel a motor response is reflected by maturational changes in the white matter underlying the right presupplementary motor area (preSMA) and posterior inferior frontal gyrus (IFG), two cortical key areas of the fronto-basal ganglia “stopping” network. Eighty-eight typically-developing children and adolescents, aged 7-19 years, were longitudinally assessed with the stop-signal task (SST) and diffusion tensor imaging (DTI) of the brain over a period of six years. Participants were examined from two to nine times with an average of 6.6 times, resulting in 576 SST-DTI datasets. We applied tract-based spatial statistics to extract mean fractional anisotropy (FA) from regions-of-interest in the white matter underlying the right IFG (IFG_FA_) and right preSMA (preSMA_FA_) at each time point. Motor response cancelation performance, estimated with the stop-signal reaction time (SSRT), improved with age. Initially well performing children plateaued around the age of 11 years, while initially poor performers caught up at the age of 13-14 years. White matter microstructure continued to mature across the investigated age range. Males generally displayed linear maturational trajectories, while females displayed more curvilinear trajectories that leveled off around 12-14 years of age. Maturational increases in right preSMA_FA_ but not right IFG_FA_ were associated with developmental improvements in SSRT. This association differed depending on the mean right preSMA_FA_ across the individual maturational trajectory. Children with lower mean right preSMA_FA_ exhibited poorer SSRT performance at younger ages but steeper developmental trajectories of SSRT improvement. Children with higher mean right preSMA_FA_ exhibited flatter trajectories of SSRT improvement along with faster SSRT already at the first assessments. The results suggest that no further improvement in motor response cancellation is achieved once a certain level of maturity in the white matter underlying the right preSMA is reached. Similar dynamics may apply to other behavioral read-outs and brain structures and, thus, need to be considered in longitudinal MRI studies designed to map brain structural correlates of behavioral changes during development.

**Highlights:**

- Motor response cancellation, i.e. SSRT, improvement plateaued at 13-14 years of age
- Fractional anisotropy (FA) captured maturation of white matter (WM) microstructure
- FA in the WM underlying right preSMA (preSMA_FA_) reflected SSRT improvement with age
- Individual SSRT improvement depended on mean right preSMA_FA_ across all DTI sessions
- Children with lower mean right preSMA_FA_ had the steepest improvements in SSRT

## 1. Introduction

The ability to effectively suppress or cancel motor response tendencies is essential for focused and goal-directed behavior. (Chambers et al., 2009; Logan, 2017). Motor response cancellation, as measured with the stop-signal task (SST), has been linked to a fronto-basal ganglia circuitry, where a primarily right-lateralized network, including the posterior inferior frontal gyrus (IFG), the presupplementary motor area (preSMA) and the subthalamic nucleus, is thought to play a key role (Aron, 2011; Chambers et al., 2009). Evidence suggests that these three regions are structurally inter-connected (Aron et al., 2007; Inase et al., 1999). While motor response cancellation has been suggested to be mediated by the IFG and preSMA through the “hyper direct” pathway, i.e. the cortico-subthalamic-pallidal pathway (Aron, 2011; Chambers et al., 2009; Nambu et al., 2002), research also suggests that the “indirect” pathway, i.e. the cortico-striatal-pallidal pathway, may play an important role in cancelling motor responses (Aron, 2011; Chambers et al., 2009; Jahfari et al., 2012; Jahfari et al., 2011). Ultimately, motor responses are inhibited by suppressing the response in the primary motor cortex and its output via the corticospinal tracts (CST).

The ability to cancel a motor response matures, i.e. stopping speed becomes faster, throughout childhood and adolescence (Curley et al., 2018; Dupuis et al., 2019; Williams et al., 1999). Developmental changes in behavior are paralleled by structural and functional maturational changes in the brain (Luna et al., 2015). Human brain structural development is a complex protracted and regionally heterogeneous process that extends well into early adulthood with higher associative cortical regions maturing later than primary sensory and motor areas (Brown et al., 2012; Jernigan et al., 2011; Tamnes et al., 2017). White matter fiber tracts, that connect brain regions and facilitate interregional communication, also exhibit variable maturational trajectories (Bava et al., 2010; Lebel et al., 2012; Lebel et al., 2008). Diffusion tensor imaging (DTI) studies in children and adolescents have consistently observed age-related increases in fractional anisotropy (FA), reflecting a disproportionate decrease in radial diffusivity (RD) relative to axial diffusivity (AD), as well as decreases in mean diffusivity (MD) across multiple white matter tracts and subcortical grey matter structures. These age-related changes are possibly due to ongoing myelination and/or increases in axonal diameter and density (Bava et al., 2010; Lebel et al., 2012; Lebel et al., 2008). The structural brain changes underlying motor response cancellation development have not been thoroughly characterized. A recent longitudinal study including 110 typically-developing children aged four to 13 years found that relatively larger cortical surface area of the right pars opercularis of the IFG was associated with faster stop-signal reaction time (SSRT), i.e. better motor response cancellation performance (Curley et al., 2018). Moreover, in a cross-sectional study of 72 typically-developing children aged seven to 13 years, we previously found that faster SSRT was associated with higher FA and lower RD in the white matter underlying both the right IFG (pars opercularis) and right preSMA (Madsen et al., 2010).

In the present longitudinal study, we set out to investigate how developmental improvements in SSRT performance were related to the maturational trajectories of the white matter microstructure within the fronto-basal ganglia “stopping” network in typically-developing children and adolescents aged 7-19 years. Based on our previous baseline findings (Madsen et al., 2010) in the same cohort, we hypothesized that individual differences in developmental improvement in SSRT would be associated with the maturational trajectories of FA and RD of the white matter underlying the right IFG and right preSMA. Additionally, we explored associations between SSRT and CST and neostriatal, i.e., caudate nucleus and putamen, microstructure. Finally, we examined for possible sex differences, because of reported sex differences in structural brain development and neural activation during the performance of the SST.

## 2. Materials and methods

### 2.1. Study design and participants

The present longitudinal study included data from 88 typically-developing children and adolescents (52 girls, 36 boys) aged 7.5-18.9 years. All participants were enrolled in the longitudinal HUBU (“*Hjernens Udvikling hos Børn og Unge*”, in English: Brain maturation in children and adolescents) project, which had been initiated in 2007, where 95 typically-developing children (55 girls, 40 boys) aged seven to 13 years and their families were recruited from three elementary schools in the Copenhagen suburban area. We included all children whose families volunteered for the HUBU project, except children who according to parent screening reports had any known history of neurological or psychiatric disorders or significant brain injury. After receiving oral and written explanation about the study aims and procedure and prior to participation, all children assented to the study procedures and informed written consent was obtained from the parents. Furthermore, we obtained written informed consent from the participants themselves when they turned 18 years of age. The study was approved by the local Danish Committee for Biomedical Research Ethics (protocols: H-KF-01-131/03 and H-3-2013-037) and conducted in accordance with the Declaration of Helsinki. Participants were assessed up to 12 times with approximately six months intervals between the first 10 assessments, and approximately one-and three-year intervals between assessments 10 and 11, and 11 and 12, respectively. Participants performed the SST at nine assessments: one through seven, nine and 11. Previous publications on cross-sectional/baseline HUBU data investigated associations between regional white and grey matter microstructure and response inhibition (Madsen et al., 2010), spatial working memory (Vestergaard et al., 2011), visuospatial choice reaction time (Madsen et al., 2011), sustained attention (Klarborg et al., 2013), circle drawing skill (Angstmann et al., 2016) and neuroticism (Madsen et al., 2018).

Seven HUBU participants (3 girls, 4 boys) were excluded entirely from the present longitudinal study because of incidental findings on MRI scans (n=1), receiving a psychiatric diagnosis after study initiation (n=2), no good quality DWI data were available (n=2), or because the participant had only been assessed at one time-point (n=2). Thus, the present study included 88 children and adolescents, and included six sibling pairs. As assessed with the Edinburg Handedness Inventory, 76 participants were right-handed, and 12 participants were left-handed. A total of 654 datasets were acquired of the 88 participants in the nine assessments that included the SST. Of these, 78 datasets were excluded because the participant did not finish the MRI scanning session (3 participants, 4 datasets), was not scanned due to metallic dental braces (11 participants, 16 datasets), had poor MR-image quality (18 participants, 23 datasets), had acquired a brain injury after baseline (1 participant, 6 datasets), did not complete the SST (1 dataset), did not perform the SST because the response box did not work (4 participants, 4 datasets) or time delays during the assessment (2 participants, 2 datasets), or because of SST performance issues due to binge drinking the night before assessment (3 participants, 3 datasets), tiredness (1 participant, 2 datasets), not understanding the task instructions (1 dataset), extremely high standard deviation (STD) on the go reaction time (1 dataset, STD = 11,115 ms) or the SSRT could not be reliably estimated (14 participants, 15 datasets, see section 2.2. Stop-signal task for further explanation). In total, 576 valid SST-MRI datasets from 88 participants were included in the present study. An overview of the longitudinal assessments and the age at each assessment for each participant are shown in Figure 1. Highest level of maternal and paternal education was acquired for all participants and translated into years of education using national norms. Average years of parental education, or years of education from one parent if unavailable for both parents, were used in the statistical analyses.

**Figure 1.**
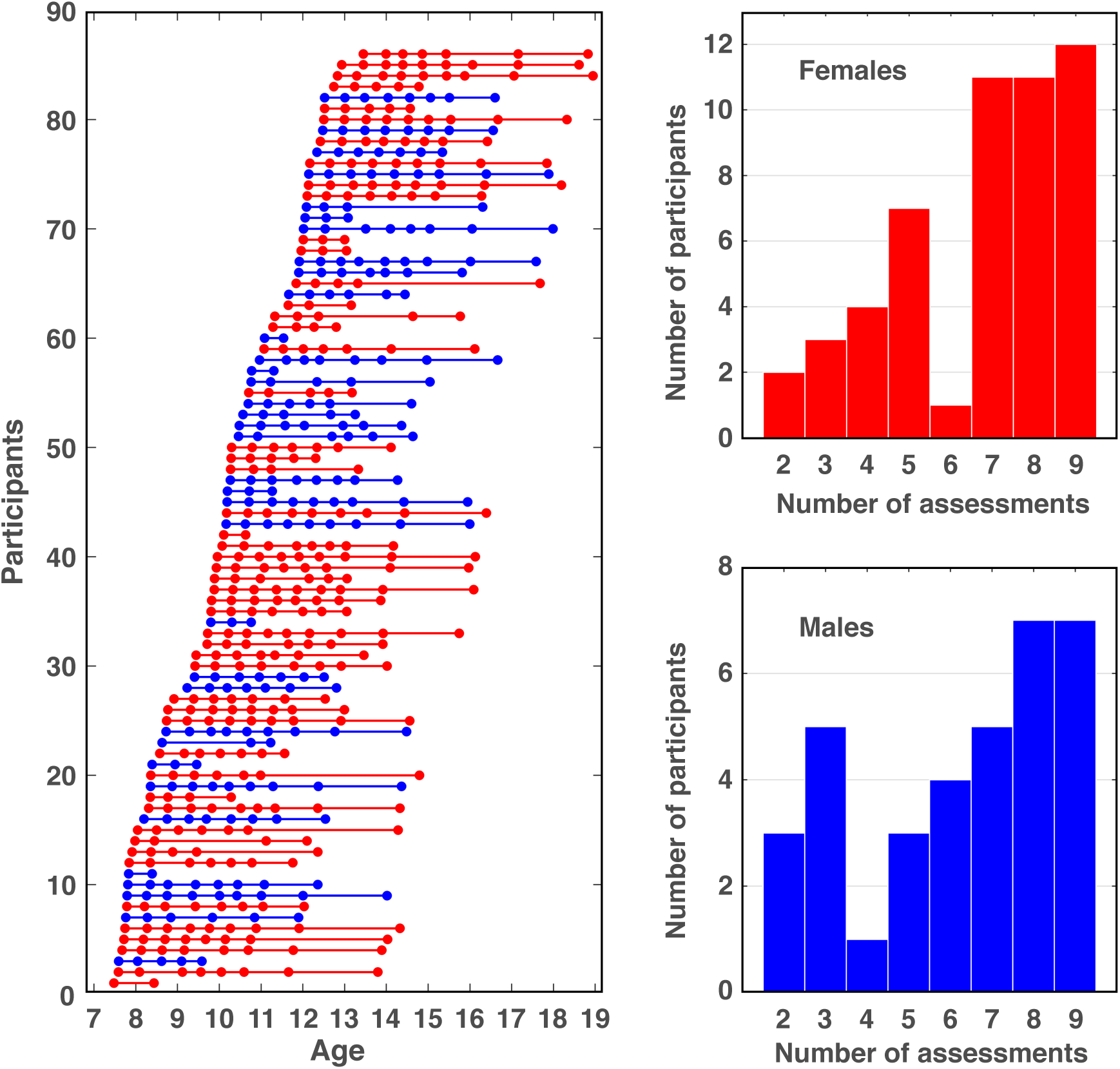
Left: Number of longitudinal assessments and age at each assessment for each of the 88 participants. Assessments are depicted by filled circles; blue for boys (n = 36) and red for girls (n = 52). In total there were 576 datasets. Right: Number of participants that have n number of assessments for boys and girls separately. Participants were assessed on average 6.6 times (range: 2 - 9) with only five participants having two timepoints.

### 2.2. Stop-signal task

“The stop-signal task (SST) was administered using the Cambridge Neuropsychological Test Automated Battery (Cambridge Cognition Ltd., Cambridge, UK). The SST consists of go and stop trials (Figure 2). Subjects sit in front of a computer monitor, with each index finger resting on a response button. A circle is presented for 500 ms, followed by an arrow pointing either left or right. Subjects are instructed to press the left or right button (i.e., with the left or right index finger), depending on the direction of the arrow, without making any mistakes, and to press as fast as possible. The test consists of two parts. In the first part, there are 16 ‘Go’ training trials without an auditory stop-signal to introduce the subjects to the press pad. In the second and longer part, an auditory stop-signal occurs in 25% of the trials. When the tone occurs, subjects must try to withhold their responses. The time between the onset of the arrow and the auditory stop signal, i.e., the stop-signal delay (SSD), changes adaptively throughout the test, depending on the subject’s past performance, so that responses are inhibited successfully approximately 50% of the time for each subject. The shorter the SSD, the more likely it is that the subject will be able to inhibit his or her response. The SST is administered in 5 blocks of 64 trials. Each block is divided into four sub-blocks of 16 trials (12 go trials and 4 stop trials in random order). There is no gap between the sub-blocks, and they are not evident to the subject. After each block, a feedback screen is displayed showing a histogram representation of the subject’s reaction time on ‘Go’ trials. The histogram shown after the first block is identical for all subjects. The test administrator explains to the subject that if he/she can go faster, it will show in the next histogram, before encouraging him/her to go faster. The feedback after each of the last four blocks is the subjects’ go reaction time in relation to the first block and consists of a histogram containing the first block and the relative performance of all previous blocks. The primary behavioral outcome measure is the stop-signal reaction time (SSRT), which measures how fast subjects can inhibit a prepotent response.” (Madsen et al., 2010). The SSRT is estimated using the race model, which assumes that the go and stop processes are in a race with each other and are (mainly) independent (Boucher et al., 2007; Logan et al., 1984). The trials in the first half of the dataset were considered training trials to familiarize subjects with the auditory stop-signal. We used the last half of the task trials to get a stable estimate of the SSD50, where subjects are able to inhibit 50% of their responses. SSRT was calculated, following the recently published consensus guidelines, using the recommended least-biased non-parametric integration method with replacement of go omissions (Verbruggen et al., 2019). Notably, there were no Go omissions in our data. If there were, the guidelines recommend replacing these with the maximum RT before calculating the SSRT. The SSRT was calculated by subtracting the mean SSD50 from the n^th^ fastest Go RT that corresponds to the proportion of trials where the subject failed to inhibit their response when a stop signal was given i.e. p(respond|signal). To determine the n^th^ fastest Go RT, RTs for all Go trials in the 20 task sub-blocks on which a response was given, irrespective of accuracy, were ordered in ascending order. The n^th^ fastest RT was than identified by multiplying the number of Go RT’s with p(respond|signal). For example, with 240 Go RTs and a p(respond|signal) of 0.40, the SSRT was calculated by subtracting the mean SSD50 from the 96^th^ fastest Go RT. Finally, a logarithm transformation was used on the SSRT (SSRT_log_) in order to stabilize the variance and used in all statistical analyses.

**Figure 2.**
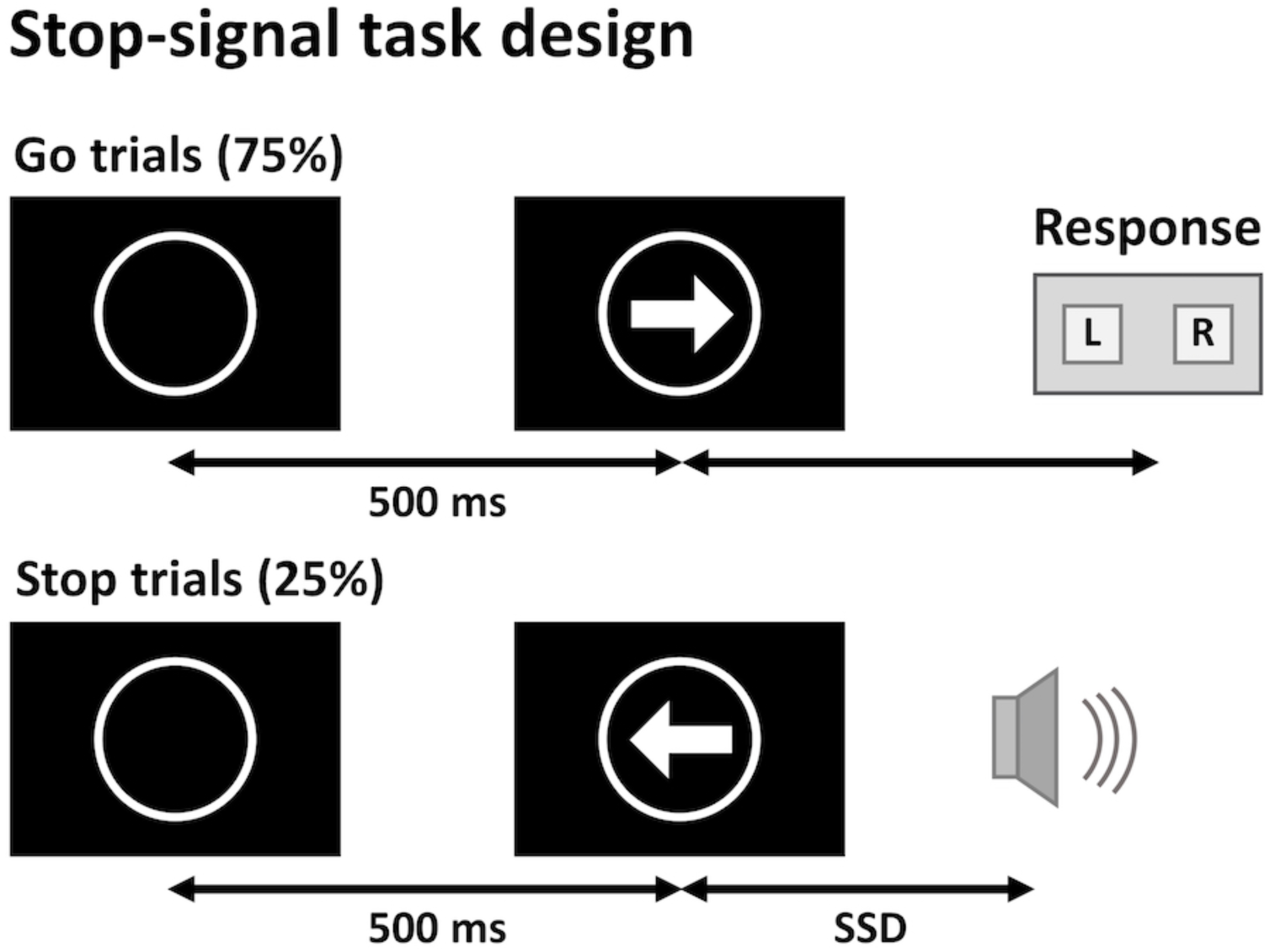
The stop-signal task consists of 75% go trial and 25% stop trials. In each trial, an empty circle is presented for 500 ms, after which an arrow pointing either to the left or the right appears inside the circle. Depending on the direction of the arrow, participants have to press the left or the right response button as fast as possible. During stop trials, after the presentation of the arrow, a sound signals the participants that they have to withhold their response. The latency to this auditory stop signal, i.e. the stop-signal delay (SSD), is dynamically varied throughout the task, in such a way that individual participants are able to inhibit approximately 50% of their responses on stop trials, i.e. the SSD50.

The SSRT could not be reliably estimated for 15 datasets from 14 participants. Of these, nine datasets from nine participants were excluded because the mean RT on unsuccessful stop trials, calculated over the last half of the task trials, were longer that the mean RT on Go trials, calculated over all Go trials with a response, irrespective of accuracy (Verbruggen et al., 2019). Six datasets from five participants were excluded because the proportion of successful stops deviated substantially from 0.5, being either lower than 0.35 or higher than 0.65. Because of the longitudinal nature of the study we chose to use thresholds that were more conservative than the recommended 0.25 and 0.75 (Verbruggen et al., 2019).

### 2.3. Image acquisition

At each assessment participants were scanned on the same “Siemens Magnetom Trio MR scanner (Siemens, Erlangen, Germany) with an eight-channel head coil (Invivo, FL, USA). All acquired scans were aligned parallel to the anterior commissure–posterior commissure line. T1-weighted images of the whole head were acquired using a 3D MPRAGE sequence (TR = 1550 ms, TE = 3.04 ms, matrix 256 x 256, 192 sagittal slices, 1 x 1 x 1 mm^3^ voxels, acquisition time = 6:38). T2-weighted images of the whole head were acquired using a 3D turbo spin echo sequence (TR = 3000 ms, TE = 354 ms, FOV = 282 x 216, matrix = 256 x 196, 192 sagittal slices, 1.1 x 1.1 x 1.1mm^3^ voxels, acquisition time = 8:29). Whole brain diffusion-weighted (DW) images were acquired using a twice-refocused balanced spin echo sequence that minimized eddy current distortion (Reese et al. 2003). Ten non-DW images (b = 0) and 61 DW images (b=1200 s/mm^2^), encoded along independent collinear diffusion gradient orientations, were acquired (TR = 8200 ms, TE = 100 ms, FOV = 220 x 220, matrix = 96 x 96, GRAPPA: factor = 2, 48 lines, 61 transverse slices with no gap, 2.3 x 2.3 x 2.3mm^3^ voxels, acquisition time = 9:50). A gradient echo field map was acquired to correct B0 field distortions (TR = 530 ms, TE[1] = 5.19 ms and TE[2] = 7.65 ms, FOV = 256 x 256; matrix = 128 x 128, 47 transverse slices with no gap, voxel size = 2 x 2 x 3 mm^3^, acquisition time = 2:18).” (Madsen et al., 2010).

### 2.4. Image evaluation

All baseline MRI scans were evaluated by an experienced neuroradiologist and all, but one, were deemed without significant clinical pathology. Prior to analysis and blind to behavioral data, all raw MR-images were visually inspected to assure sufficient image quality. Based on this inspection, 25 datasets were excluded from further analyses due to poor image quality (see section “Study design and participants”).

### 2.5. Construction of the diffusion tensor images

Image preprocessing was done using MATLAB scripts that were mainly based on SPM 8 routines (Wellcome Department of Cognitive Neurology, University College London, UK). The T1-and T2-weighted images were rigidly oriented to MNI space (six-parameter mutual information) and corrected for spatial distortions due to nonlinearity in the gradient system of the scanner (Jovicich et al., 2006) (note that at this point no-reslicing was performed). T2-weighted images were then rigidly coregistered to the T1-weighted image. To align the DWI images to the T1-weighted image, the mean b0 image was first rigidly registered to the T2-weighted image, after which all DW images were coregistered to the mean b0 image (no reslicing). Next, all co-registered DWI images were corrected for spatial distortions using a voxel displacement map based on the acquired b0 field map (Andersson et al., 2001) and the scanner-specific gradient non-linearity profile (Jovicich et al., 2006). Subsequently, all images were resliced using tri-linear interpolation. Importantly, the above procedure ensures that only one re-slicing step was employed. Diffusion gradient orientations were adjusted to account for any applied rotations. The least-squares-fit by non-linear optimization, employing a Levenburg-Marquardt algorithm and constrained to be positive definite by fitting its Cholesky decomposition, implemented in Camino was used to fit the diffusion tensor (DT) (Jones and Basser, 2004). Finally, for each subject we estimated movement during DWI scanning by calculating the root mean square deviation of the six rigid body transformation parameters resulting from coregistering DWI images to the mean b0 image (Jenkinson 1999, Taylor et al., 2016), using a spherical volume radius of 60 mm to approximate the brain.

### 2.6. Spatial normalization of the longitudinal diffusion tensor images

The DT images were spatially normalized using DTI-TK, which uses the high dimensional information of the diffusion tensor to achieve highly accurate normalizations (Zhang et al., 2007). We employed an unbiased longitudinal two-step approach (Keihaninejad et al., 2013). First, within each subject all DT images over all time points were registered together to create within-subject DT image templates. Secondly, within-subject template DT images were registered together to create a between-subject DT template image. Concatenation of the within- and between subject registration deformation fields was used to warp individual DWI volumes into a common study specify space. Fractional anisotropy (FA), mean diffusivity (MD), axial diffusivity (AD = λ1) and radial diffusivity (RD = (λ2 + λ3) / 2) images were created. Finally, non-brain voxels in FA and diffusivity images were removed by employing a brain mask based on warped b0 images.

### 2.7. Tract-based spatial statistics

Tract-Based Spatial Statistics (TBSS) (Smith et al. 2006), part of FSL 5.0.9, was used to create a mean FA skeleton, representing the centers of all tracts common to the group. Instead of using the standard TBSS normalization steps, the between-subject FA template image (from DTI-TK) was aligned to MNI space using affine registration (flirt, FSL). Subsequently, the between-subject diffusivity images and the normalized individual DT images were transformed into 1 mm^3^ MNI space. Next, the MNI space aligned, between-subject FA template image was entered into the TBSS processing stream using the “tbss_skeleton” script, part of the “tbss_3_postreg” processing step, in which the between-subject FA template image was thinned to create a mean FA skeleton. The mean FA skeleton was thresholded at FA > 0.2 and contained 102.983 1mm^3^ interpolated isotopic voxels, corresponding to approximately 22% of the voxels (in the mean FA map across subjects and time points) with FA above 0.2. All participants’ aligned FA images were then projected onto the mean FA skeleton by locating the voxels with the highest local FA value perpendicular to the skeleton tracts and assigning these values to the skeleton. Finally, the skeleton projections were applied on the MD, AD and RD data.

### 2.8. Spatial normalization of grey matter

The MNI-oriented T1 and T2-weighted images were processed using the VBM8 toolbox in SPM8 (http://dbm.neuro.uni-jena.de/vbm8/VBM8-Manual.pdf), which includes a unified segmentation algorithm (Ashburner and Friston, 2005) and a Hidden Markov Random Field (HMRF) method (Cuadra et al., 2005). T2-weighted images were used to automatically create brain masks. Resulting brain masked grey and white matter tissue maps in native space, and the affine part of the spatial transformation from native to MNI space (obtained from the VBM8 analysis) were subsequently used in DARTEL (“Diffeomorphic Anatomical Registration Through Exponentiated Lie Algebra”), using default settings (Ashburner, 2007) for high-dimensional intra-subject and inter-subject registration. Average inter-subject and between-subject T1-weighted templates were generated and used for drawing regions-of-interest (ROIs) in the subcortical grey matter structures.

### 2.9. Regions-of-interest (ROIs)

In the present study, we extracted FA, MD, RD and AD values from white and grey matter ROIs to test specific hypotheses and to determine the anatomical specificity of observed associations (see below).

#### 2.9.1. White matter ROIs

White matter ROIs were drawn manually onto the mean skeleton in the white matter underlying the left and right inferior frontal gyrus (IFG, pars opercularis) and pre-supplementary motor area (preSMA) as well as in the corticospinal tracts (CST) overlaid on the mean FA image using FSLview. The ROIs are shown in Figure 3.

**Figure 3.**
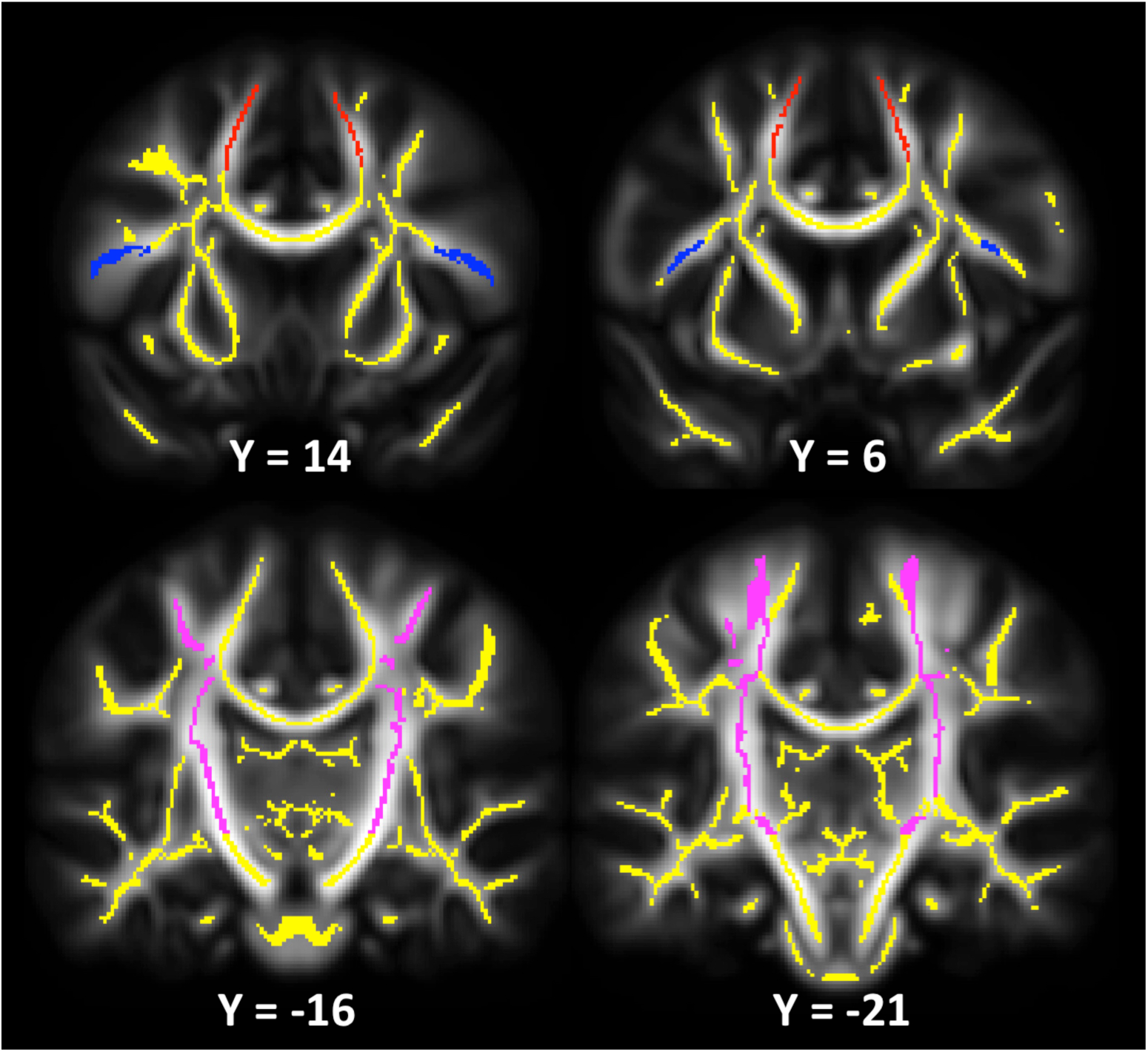
Coronal views of the regions-of-interest in the white matter underlying the inferior frontal gyrus (pars opercularis, blue) and the presupplementary motor area (preSMA, red) as well as the corticospinal tracts (magenta) overlaid on the TBSS skeleton (yellow) and the mean fractional anisotropy (FA) map. The corresponding MNI coordinates (Y) are given below each slice.

“The right and left pars opercularis were located using a brain atlas (Duvernoy, 1999) and published information about the morphology of the pars opercularis (Tomaiuolo et al., 1999). The boundaries of the IFG ROIs were found by using anatomical information visible in the target FA map. The vertical ramus of the lateral fissure and the inferior part of the precentral sulcus, indicated by low FA values, were used as landmarks to define the anterior and posterior boundaries of the pars opercularis. The skeleton defined the superior boundary, in that the lateral part of the pars opercularis segment was perpendicular to the segment going into the pars triangularis.” (Madsen et al., 2010). The right IFG ROI contained 348 voxels and the left contained 339 voxels.

The preSMA ROIs were located using MNI coordinates derived from previous structural and functional studies (Behrens et al., 2006; Johansen-Berg et al., 2004) and defined between the MNI coordinates y = 0 and y = 20. However, to ensure that the ROIs mainly included white matter skeleton voxels extending towards the preSMA, the posterior and anterior boundaries were set to MNI y= 5 and y = 15 in the present study. The right preSMA ROI included 287 voxels and the left ROI included 326 voxels.

The CST ROIs included skeleton segments extending from the motor cortex (hand area) and the sensorimotor cortex, and down through the posterior limb of the internal capsule. Below the level of the corpus callosum, the probabilistic fiber atlas implemented in FSLview (Hua et al., 2008) was used to guide the delineation of the CST ROIs. The inferior boundary for the CST ROIs was set at the level of the anterior commissure (z = −5). Furthermore, the ROIs were located between the slices y = −12 to y = −38. The right and left CST ROIs contained 1.921 and 2.024 voxels, respectively.

Finally, mean FA, MD, RD and AD values from the left and right ROIs as well as from the whole skeleton were extracted from each of the participants’ scans for statistical analyses.

#### 2.9.2. Grey matter ROIs

ROIs of the caudate nuclei and putamen were constructed to extract MD of these structures from each participant. ROIs were drawn onto the DARTEL-warped between-subject T1-weighted template image (Figure 4). Striatal cell bridges were excluded from any of the ROIs. The posterior border of the caudate nucleus ROI was defined by the posterior commissure in the coronal plane, and, thus, the posterior part of the tail of the caudate nucleus was excluded. Furthermore, we constructed an ROI of nearby, but functionally distinct, subcortical grey matter, namely the combined nucleus accumbens and amygdala, in order to examine the anatomical specificity of possible neostriatal effects. This ROI, thus, served as a control region for comparison to the neostriatal ROIs. Several steps were conducted to ensure that the subcortical ROIs, for which MD was extracted, only included grey matter voxels by excluding partial-volume voxels. First, the between-subject FA and RD template images were aligned to the DARTEL template using affine registration. The binary ROIs were then multiplied with the between-subject template FA image and thresholded at FA<0.3 to exclude potential white matter partial-volume voxels, and subsequently multiplied with the between-subject RD template image and thresholded at RD < 0.0012 (10^-3^ mm^2^/s) to exclude voxels likely to include cerebrospinal fluid. Second, the ROIs were eroded to remove an approximately one-voxel thick layer of the outer border around the ROIs (Figure 4). Third, the resulting thresholded and eroded ROIs were aligned to DW template space using the inverted affine registration matrix, where the binary ROIs were multiplied with the participants’ own (intra-subject) FA and RD template images and thresholded as above. Lastly, the final ROIs were overlaid on the participants’ FA template image to visually inspect the anatomical fit of the ROIs for each participant. Based on the visual inspection, the caudate nucleus was excluded for one participant due to poor fit. The mean (± std) volume (mm^3^) of the ROIs were: left putamen = 3,427 (± 1), right putamen = 3,535 (± 0), left caudate nucleus = 2,375 (± 148), and right caudate nucleus = 2,611 (± 146).

**Figure 4.**
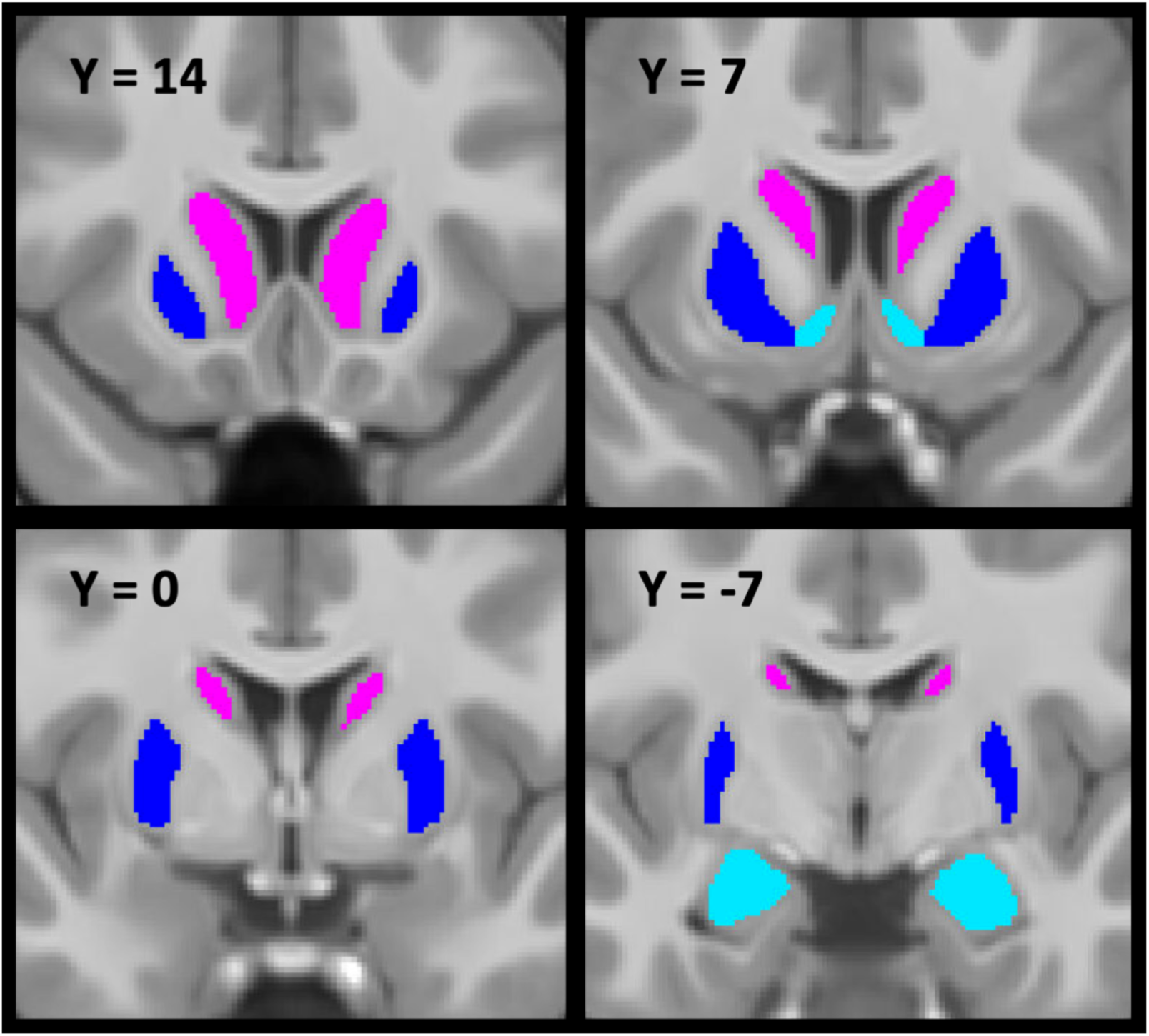
Coronal views of the thresholded and eroded region-of-interest (ROIs). The putamina in blue, the caudate nuclei in magenta and the nuclei accumbens/amygdalae control regions in light blue are overlaid on the DARTEL-warped between-subject T1-weighted template. The corresponding MNI coordinates (Y) are given for each slice.

### 2.10. Statistical analyses

All statistical analyses were performed in R version 3.6.1 (R Core Team, 2019).

#### 2.10.1. Age, sex and age-by-sex effects on outcome measures of interest

Before testing our hypotheses, we used generalized additive mixed models (GAMM), using the mgcv package, version 1.8-29 and the nlme package, version 3.1-141, to characterize possible age, sex and age-by-sex interaction effects on SSRT, SSRT_log_, median go RT (Go RT_median_), Go RT_STD_, SSD50, left and right IFG FA (IFG_FA_) and RD (IFG_RD_), and preSMA FA (PreSMA_FA_) and RD (PreSMA_RD_), total CST and neostriatal MD, left and right CST MD, FA and RD as well as left and right caudate nucleus and putamen MD. GAMM uses non-linear smooth functions to fit the complex longitudinal data. The “estimated degrees of freedom” (edf) reflect the degrees of freedom that are used by a smooth function to fit the data, with an edf of one representing a straight line or a linear relationship. The following generic model was tested: dependent measure (i.e. SSRT or ROI DTI) = s(age) + sex + ti(age, by = sex). Ordered sex (female = 1, male = 0) was used as a covariate, and the smooth terms “s” and “ti” were used for respectively age and age-by-sex interaction effects, while participant and within-subject age [random = list (participant id = ∼1 + age)] were used as random effects to respectively fit individual intercepts and slopes. To fit the data, we used the default thin plate regression splines smoothing function with default penalizing and five knots (k). k represents the k - 1 number of basis functions used to generate the smooth function, which was set to five since k>5, in some cases, led to unrealistically “wavy” or in GAMM terms “wiggly” curves, indicative of overfitting. Finally, the smoothing parameters were estimated using the restricted maximum likelihood (REML) method. Continuous variables i.e. age, SST variables and ROI DTI measures were converted to z-scores.

#### 2.10.2. Testing for possible test-retest effects on SSRT

Given the longitudinal nature of the SST data, we examined test-retest (including practice/learning) effects on the SSRT. Test-retest effects may be present if participants performing the SST for the second time perform significantly better (i.e. lower SSRT) than individuals who were of the same age when performing the SST for the first time. Because participants’ ages increase with each SST assessment round, we conducted a GAMM as described above to adjust SSRT for any linear or non-linear age, sex and age-by-sex effects, and extracted the SSRT residuals. Next, we used the SSRT residuals in paired t-tests to test for possible differences in SSRT between adjacent assessments for the first four SST assessments. In addition, we tested if the test-retest effect differed with age. To this end, we subtracted SSRT from adjacent assessment rounds (e.g. SSRT round 1 – SSRT round 2) and used each change score as the dependent variable in three separate linear regression models, with participants’ mean age across the two rounds as the independent variable.

#### 2.10.3. Individual differences in the developmental trajectories of SSRT

We examined if initially good and poor performers differed in their developmental trajectories of SSRT performance. First, we ranked the GAMM SSRT residuals from the first and second SST assessments separately and averaged these to get a stable estimate of the participantś initial performance. The average ranking was then used to split the participants into three groups: good (n=30), intermediate (n=30) and poor (n=28) performers. To visualize group-specific developmental trajectories, we conducted a GAMM for each group, where SSRT was fitted with s(age) with random effects for participant and within-subject age to fit individual intercepts and slopes. To test if the groups differed in their developmental trajectories, we constructed the following linear mixed effects (LME) model using the nlme package, version 3.1-141: SSRT_log_ = age + age^2^ + group + age*group + age^2^*group. Participant and within-subject age were entered as random effects.

#### 2.10.4. Testing a priori hypotheses

To test our a priori hypotheses, we used LME models. We hypothesized that developmental changes in SSRT performance would be associated with individual differences in the maturational trajectories of FA and RD of the white matter underlying right IFG and right preSMA, after controlling for age effects. Since both response cancellation performance and white matter microstructure change with age, we included age as a covariate to account for any non-specific association attributable to unmeasured factors indexed by chronological age. The exact tested models depended on the results of the initial GAMM analyses (see Table 2, results section). Because of a nonlinear effect of age on SSRT_log_, age^2^ was entered as an additional covariate in all models. Moreover, because the age effects were linear for the four a priori ROI microstructural measures, the a priori models only included an age-by-ROI microstructure term and no age^2^-by-ROI microstructure term. Furthermore, based on observing significant linear and non-linear age-by-sex effects, sex and sex-by-age interaction terms were added respectively to right IFG_FA_, and right IFG_RD_ and preSMA_RD_ models. Additionally, since we observed a significant test-retest effect for SSRT between the first and second SST round (see section 3.1.), we included SST_test-retest_ (baseline = 0, follow-ups = 1) as a covariate in all models. Participant and within-subject age were entered as random effects. The following models were used to test our hypotheses.

For right IFG_FA_: Model 1

SSRT_log_ = SST_test-retest_ + age + age^2^ + sex + age*sex + ROI + age*ROI.

For right PreSMA_FA_: Model 2

SSRT_log_ = SST_test-retest_ + age + age^2^ + ROI + age*ROI

For right IFG_RD_ and right PreSMA_RD_: Models 3 and 4

SSRT_log_ = SST_test-retest_ + age + age^2^ + sex + age*sex + age^2^*sex + ROI + age*ROI

In testing our a priori hypotheses, we were only interested in the age-by-ROI microstructure interactions. We therefore used a Bonferroni-corrected p-value of 0.05/4 (e.g. one age-by-ROI interaction term for each of the ROI FA and RD measures) = 0.0125 to correct for multiple comparisons. Contingent on any observed effects, we conducted several follow-up models to determine if effects remained when correcting for average years of parent education, handedness and movement during DWI scanning. Furthermore, we tested for possible anatomical specificity of any observed effects by adjusting for either whole brain or left hemisphere ROI DTI measures. Finally, to obtain more information about the nature of possible FA findings, we examined ROI AD, as increases in FA could be due to decreases in RD and/or increases in AD. All follow-up models were tested at the α = 0.05 level.

#### 2.10.5. Exploratory analyses

We investigated possible associations between SSRT_log_ and CST and neostriatal MD using the same modeling strategy as when testing our a priori hypotheses. A combined nucleus accumbens and amygdala gray matter microstructure measure was used to examine the anatomical specificity of any observed neostriatal effects. Based on the initial GAMM analyses (see Table 2, and section 3. Results), we tested the following model: SSRT_log_ = SST_test-retest +_ age + age^2^ + sex + age*sex+ age^2^*sex+ ROI + age*ROI + age^2^*ROI. In our exploratory analyses we did not correct for multiple comparisons and considered a p < 0.05 for any of the age-by-ROI or age^2^-by-ROI interaction effects to be of interest.

#### 2.10.5. Whole TBSS skeleton effect size maps

Finally, we created an effect size map to provide further information about the association between SSRT_log_ and age-by-FA across the white matter skeleton. The effect size map is an unthresholded t-map of the association between SSRT_log_ and age-by-FA that was generated using the following LME model: SSRT_log_ = SST_test-retest +_ DWI movement + age + age^2^ + sex + age*sex + age^2^*sex + FA + **age*FA**. The unthresholded t-map has been uploaded to NeuroVault.org (Gorgolewski et al., 2015) and is available at https://neurovault.org/collections/AMYMNIEX/.

## 3. Results

### 3.1. Age, sex, age-by-sex and test-retest effects on the SST measures

SST measures from the nine assessment rounds are presented in Table 1. The proportion of successful stops was on average approximately 0.5 in all assessment rounds, indicating that the dynamic tracking algorithm performed as expected. None of the participants committed any go omissions. For SSRT_log_, there was one outlier that was more than four standard deviations below the mean. Since there were no objective reasons to exclude this data point and because running our analyses with and without it did not change any of our results, we did not exclude the outlier from any of our plots or analyses. We observed a significant difference in age and sex adjusted SSRT residuals between the first and second time the participants performed the SST (df = 75, t = 3.212, p = 0.0019), but no significant differences between the second and third (df = 74, t = 1.113, p = 0.269), or between the third and fourth (df = 65, t = 0.355, p =0.724) SST assessment (see Figure 5G). Moreover, we did not observe any significant differences in test-retest effects with age (p’s > 0.268), suggesting that test-retest effects did not differ for younger and older children. Since we only observed test-retest effects between the first and second SST assessment, we included a dummy variable (SST_test-retest_: baseline = 0, follow-ups = 1) in all analyses including SST variables to account for the test-retest effect.

**Figure 5.**
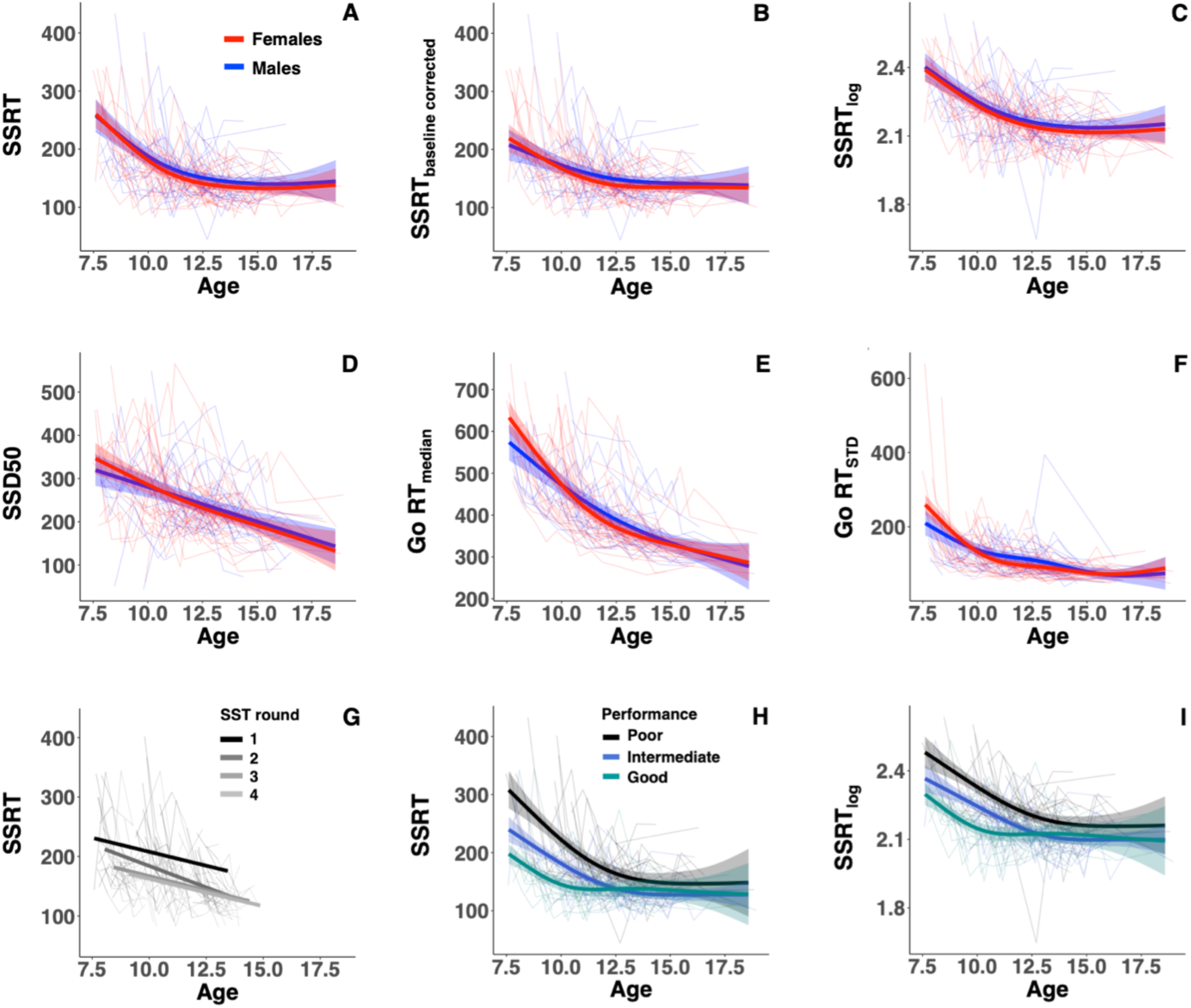
Spaghetti plots overlaid with GAMM estimated age trajectories with shaded 95% confidence intervals for males (blue) and females (red) for stop-signal reaction time (SSRT, A), SSRT corrected for test-retest effects between baseline and follow-up assessments (B) logarithm-transformed SSRT (SSRT_log_, C), stop-signal delay 50% (SSD50, D), Median Go reaction time (E), and standard deviation on Go reaction time (F). (G) displays linear regression lines for SSRT as a function of age for each of the first four SST assessment rounds separately. SSRT significantly differed between rounds 1 and 2, visualized by the jump between the lines for round 1 and 2. (H) and (I) display spaghetti plots overlaid with GAMM estimated age trajectories with shaded 95% confidence intervals for initial poor (black), intermediate (blue) and good (cyan) performers for respectively SSRT and SSRT_log_. Faster SSRT and SSRT_log_ denote better motor response cancelation performance.

**Table 1.**
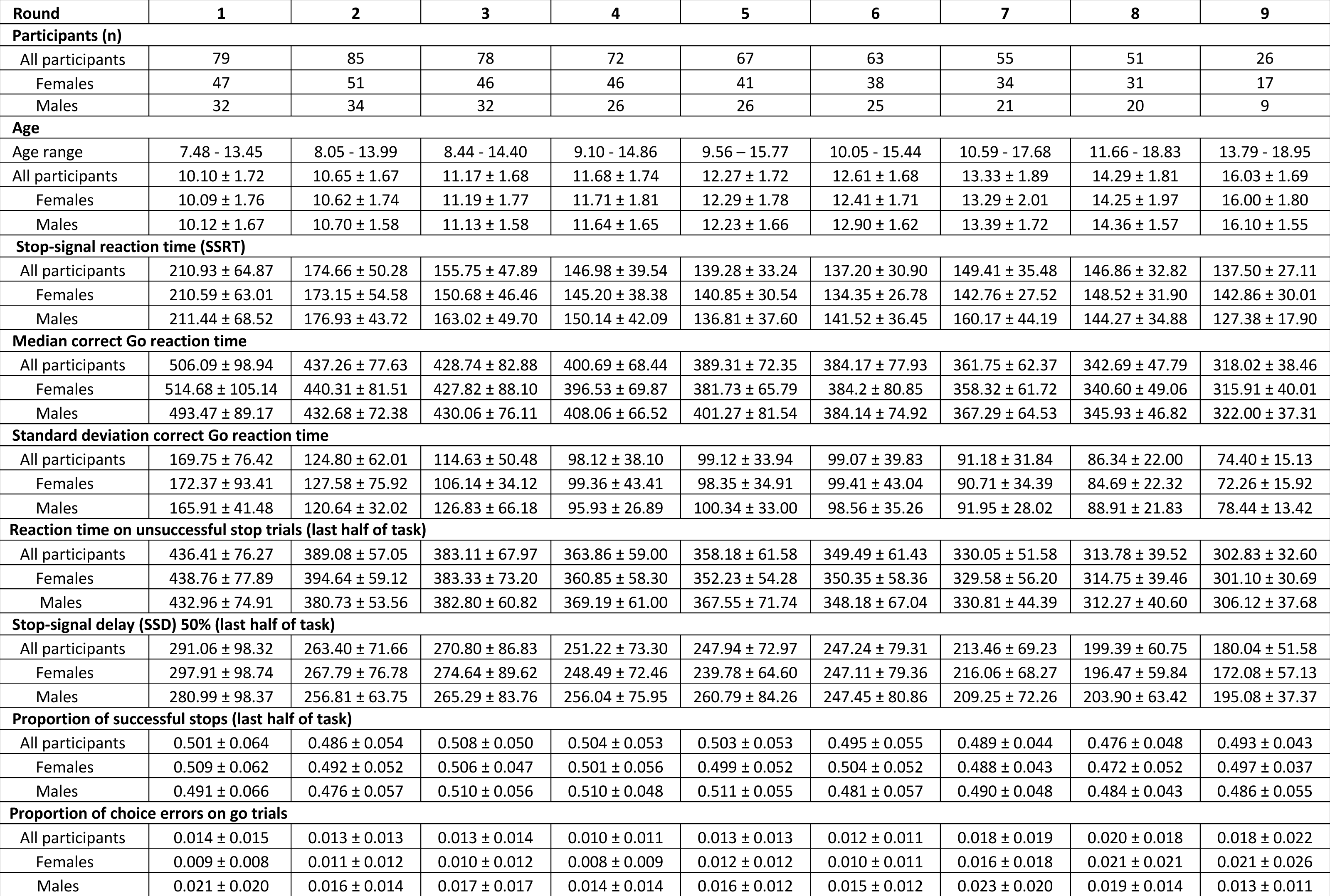
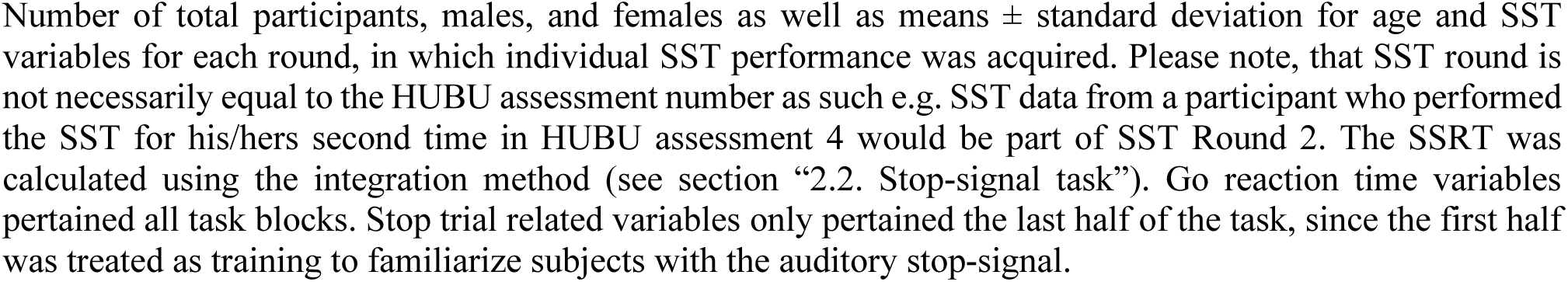
Summary of the stop-signal task (SST) variables across SST assessment rounds

| Round | 1 | 2 | 3 | 4 | 5 | 6 | 7 | 8 | 9 |
| --- | --- | --- | --- | --- | --- | --- | --- | --- | --- |
| **Participants (n)** |  |  |  |  |  |  |  |  |  |
| All participants | 79 | 85 | 78 | 72 | 67 | 63 | 55 | 51 | 26 |
| Females | 47 | 51 | 46 | 46 | 41 | 38 | 34 | 31 | 17 |
| Males | 32 | 34 | 32 | 26 | 26 | 25 | 21 | 20 | 9 |
| **Age** |  |  |  |  |  |  |  |  |  |
| Age range | 7.48 - 13.45 | 8.05 - 13.99 | 8.44 - 14.40 | 9.10 - 14.86 | 9.56 - 15.77 | 10.05 - 15.44 | 10.59 - 17.68 | 11.66 - 18.83 | 13.79 - 18.95 |
| All participants | 10.10 ± 1.72 | 10.65 ± 1.67 | 11.17 ± 1.68 | 11.68 ± 1.74 | 12.27 ± 1.72 | 12.61 ± 1.68 | 13.33 ± 1.89 | 14.29 ± 1.81 | 16.03 ± 1.69 |
| Females | 10.09 ± 1.76 | 10.62 ± 1.74 | 11.19 ± 1.77 | 11.71 ± 1.81 | 12.29 ± 1.78 | 12.41 ± 1.71 | 13.29 ± 2.01 | 14.25 ± 1.97 | 16.00 ± 1.80 |
| Males | 10.12 ± 1.67 | 10.70 ± 1.58 | 11.13 ± 1.58 | 11.64 ± 1.65 | 12.23 ± 1.66 | 12.90 ± 1.62 | 13.39 ± 1.72 | 14.36 ± 1.57 | 16.10 ± 1.55 |
| **Stop-signal reaction time (SSRT)** |  |  |  |  |  |  |  |  |  |
| All participants | 210.93 ± 64.87 | 174.66 ± 50.28 | 155.75 ± 47.89 | 146.98 ± 39.54 | 139.28 ± 33.24 | 137.20 ± 30.90 | 149.41 ± 35.48 | 146.86 ± 32.82 | 137.50 ± 27.11 |
| Females | 210.59 ± 63.01 | 173.15 ± 54.58 | 150.68 ± 46.46 | 145.20 ± 38.38 | 140.85 ± 30.54 | 134.35 ± 26.78 | 142.76 ± 27.52 | 148.52 ± 31.90 | 142.86 ± 30.01 |
| Males | 211.44 ± 68.52 | 176.93 ± 43.72 | 163.02 ± 49.70 | 150.14 ± 42.09 | 136.81 ± 37.60 | 141.52 ± 36.45 | 160.17 ± 44.19 | 144.27 ± 34.88 | 127.38 ± 17.90 |
| **Median correct Go reaction time** |  |  |  |  |  |  |  |  |  |
| All participants | 506.09 ± 98.94 | 437.26 ± 77.63 | 428.74 ± 82.88 | 400.69 ± 68.44 | 389.31 ± 72.35 | 384.17 ± 77.93 | 361.75 ± 62.37 | 342.69 ± 47.79 | 318.02 ± 38.46 |
| Females | 514.68 ± 105.14 | 440.31 ± 81.51 | 427.82 ± 88.10 | 396.53 ± 69.87 | 381.73 ± 65.79 | 384.2 ± 80.85 | 358.32 ± 61.72 | 340.60 ± 49.06 | 315.91 ± 40.01 |
| Males | 493.47 ± 89.17 | 432.68 ± 72.38 | 430.06 ± 76.11 | 408.06 ± 66.52 | 401.27 ± 81.54 | 384.14 ± 74.92 | 367.29 ± 64.53 | 345.93 ± 46.82 | 322.00 ± 37.31 |
| **Standard deviation correct Go reaction time** |  |  |  |  |  |  |  |  |  |
| All participants | 169.75 ± 76.42 | 124.80 ± 62.01 | 114.63 ± 50.48 | 98.12 ± 38.10 | 99.12 ± 33.94 | 99.07 ± 39.83 | 91.18 ± 31.84 | 86.34 ± 22.00 | 74.40 ± 15.13 |
| Females | 172.37 ± 93.41 | 127.58 ± 75.92 | 106.14 ± 34.12 | 99.36 ± 43.41 | 98.35 ± 34.91 | 99.41 ± 43.04 | 90.71 ± 34.39 | 84.69 ± 22.32 | 72.26 ± 15.92 |
| Males | 165.91 ± 41.48 | 120.64 ± 32.02 | 126.83 ± 66.18 | 95.93 ± 26.89 | 100.34 ± 33.00 | 98.56 ± 35.26 | 91.95 ± 28.02 | 88.91 ± 21.83 | 78.44 ± 13.42 |
| **Reaction time on unsuccessful stop trials (last half of task)** |  |  |  |  |  |  |  |  |  |
| All participants | 436.41 ± 76.27 | 389.08 ± 57.05 | 383.11 ± 67.97 | 363.86 ± 59.00 | 358.18 ± 61.58 | 349.49 ± 61.43 | 330.05 ± 51.58 | 313.78 ± 39.52 | 302.83 ± 32.60 |
| Females | 438.76 ± 77.89 | 394.64 ± 59.12 | 383.33 ± 73.20 | 360.85 ± 58.30 | 352.23 ± 54.28 | 350.35 ± 58.36 | 329.58 ± 56.20 | 314.75 ± 39.46 | 301.10 ± 30.69 |
| Males | 432.96 ± 74.91 | 380.73 ± 53.56 | 382.80 ± 60.82 | 369.19 ± 61.00 | 367.55 ± 71.74 | 348.18 ± 67.04 | 330.81 ± 44.39 | 312.27 ± 40.60 | 306.12 ± 37.68 |
| **Stop-signal delay (SSD) 50% (last half of task)** |  |  |  |  |  |  |  |  |  |
| All participants | 291.06 ± 98.32 | 263.40 ± 71.66 | 270.80 ± 86.83 | 251.22 ± 73.30 | 247.94 ± 72.97 | 247.24 ± 79.31 | 213.46 ± 69.23 | 199.39 ± 60.75 | 180.04 ± 51.58 |
| Females | 297.91 ± 98.74 | 267.79 ± 76.78 | 274.64 ± 89.62 | 248.49 ± 72.46 | 239.78 ± 64.60 | 247.11 ± 79.36 | 216.06 ± 68.27 | 196.47 ± 59.84 | 172.08 ± 57.13 |
| Males | 280.99 ± 98.37 | 256.81 ± 63.75 | 265.29 ± 83.76 | 256.04 ± 75.95 | 260.79 ± 84.26 | 247.45 ± 80.86 | 209.25 ± 72.26 | 203.90 ± 63.42 | 195.08 ± 37.37 |
| **Proportion of successful stops (last half of task)** |  |  |  |  |  |  |  |  |  |
| All participants | 0.501 ± 0.064 | 0.486 ± 0.054 | 0.508 ± 0.050 | 0.504 ± 0.053 | 0.503 ± 0.053 | 0.495 ± 0.055 | 0.489 ± 0.044 | 0.476 ± 0.048 | 0.493 ± 0.043 |
| Females | 0.509 ± 0.062 | 0.492 ± 0.052 | 0.506 ± 0.047 | 0.501 ± 0.056 | 0.499 ± 0.052 | 0.504 ± 0.052 | 0.488 ± 0.043 | 0.472 ± 0.052 | 0.497 ± 0.037 |
| Males | 0.491 ± 0.066 | 0.476 ± 0.057 | 0.510 ± 0.056 | 0.510 ± 0.048 | 0.511 ± 0.055 | 0.481 ± 0.057 | 0.490 ± 0.048 | 0.484 ± 0.043 | 0.486 ± 0.055 |
| **Proportion of choice errors on go trials** |  |  |  |  |  |  |  |  |  |
| All participants | 0.014 ± 0.015 | 0.013 ± 0.013 | 0.013 ± 0.014 | 0.010 ± 0.011 | 0.013 ± 0.013 | 0.012 ± 0.011 | 0.018 ± 0.019 | 0.020 ± 0.018 | 0.018 ± 0.022 |
| Females | 0.009 ± 0.008 | 0.011 ± 0.012 | 0.010 ± 0.012 | 0.008 ± 0.009 | 0.012 ± 0.012 | 0.010 ± 0.011 | 0.016 ± 0.018 | 0.021 ± 0.021 | 0.021 ± 0.026 |
| Males | 0.021 ± 0.020 | 0.016 ± 0.014 | 0.017 ± 0.017 | 0.014 ± 0.014 | 0.016 ± 0.012 | 0.015 ± 0.012 | 0.023 ± 0.020 | 0.019 ± 0.014 | 0.013 ± 0.011 |

Number of total participants, males, and females as well as means ± standard deviation for age and SST variables for each round, in which individual SST performance was acquired. Please note, that SST round is not necessarily equal to the HUBU assessment number as such e.g. SST data from a participant who performed the SST for his/hers second time in HUBU assessment 4 would be part of SST Round 2. The SSRT was calculated using the integration method (see section “2.2. Stop-signal task”). Go reaction time variables pertained all task blocks. Stop trial related variables only pertained the last half of the task, since the first half was treated as training to familiarize subjects with the auditory stop-signal.

Effects of age, sex, age-by-sex and SSRT_test-retest_ on the SST behavioral measures are reported in Table 2. Spaghetti plots overlaid with GAMM estimated age trajectories for males and females are shown in Figure 5A-F. The SSRT_log_ became significantly faster with age, suggesting that the participants became better at motor response cancellation as they grew older, which leveled off around 13-14 years of age. Median correct go RT (Go RT_median_) became faster and the standard deviation for the go RT (Go RT_STD_) became smaller with age, suggesting the participants became faster and less variable in their motor responses with increasing age. Additionally, the SSD50 became significantly shorter with age. We did not observe significant sex or age-by-sex interaction effects for any of the SST behavioral measures, except for a significant age-by-sex effect for the Go RT_median_ and Go RT_STD_. For these, males displayed a more linear decline with age, whereas females showed a curvilinear decline that leveled off earlier than in males. The SST_test-retest_ variable was significantly associated with SSRT, SSRT_log_, Go RT_median_, and Go RT_STD_, but not with SSD50 (see legend of Table 2).

**Table 2.** GAMM models: age, sex and age-by-sex effects on SST and ROI DTI measures

|  | Age |  |  | Sex |  | Age-by-sex |  |  |
| --- | --- | --- | --- | --- | --- | --- | --- | --- |
| Measure | edf | F | p | t | p | edf | F | p |
| SSRT | 2.725 | 8.306 | $< 10^{-4}$ | -1.110 | 0.267 | 2.273 | 0.528 | 0.437 |
| SSRT<sub>log</sub> | 2.827 | 8.643 | $< 10^{-4}$ | -1.016 | 0.310 | 1.654 | 0.443 | 0.684 |
| SSD50 | 1.014 | 21.364 | $< 10^{-5}$ | -0.211 | 0.833 | 1.818 | 1.173 | 0.404 |
| Go RT<sub>median</sub> | 1.830 | 40.896 | $< 10^{-14}$ | -0.301 | 0.764 | 3.249 | 5.722 | **0.0004** |
| Go RT<sub>STD</sub> | 1.001 | 25.82 | $< 10^{-6}$ | -0.726 | 0.468 | 3.714 | 15.90 | $< 10^{-9}$ |
| Right IFG FA | 1.007 | 248.385 | $< 10^{-15}$ | -0.950 | 0.343 | 1.000 | 8.049 | **0.005** |
| Right IFG RD | 1.008 | 332.443 | $< 10^{-15}$ | -0.095 | 0.925 | 2.311 | 9.246 | $< 10^{-4}$ |
| Right preSMA FA | 1.088 | 101.472 | $< 10^{-15}$ | -0.599 | 0.549 | 1.932 | 0.944 | 0.361 |
| Right preSMA RD | 1.084 | 270.494 | $< 10^{-15}$ | -0.118 | 0.906 | 2.877 | 7.797 | $< 10^{-4}$ |
| CST MD | 1.943 | 120.786 | $< 10^{-15}$ | 0.561 | 0.5765 | 2.948 | 6.355 | **0.0004** |
| Striatum MD | 2.454 | 17.629 | $< 10^{-8}$ | -1.374 | 0.170 | 3.033 | 5.145 | **0.002** |
| Left IFG FA | 1.007 | 28.142 | $< 10^{-6}$ | 0.621 | 0.535 | 1.000 | 0.954 | 0.329 |
| Left IFG RD | 2.068 | 42.341 | $< 10^{-15}$ | -0.819 | 0.413 | 2.338 | 2.324 | 0.07 |
| Left preSMA FA | 2.801 | 49.06 | $< 10^{-15}$ | -0.166 | 0.868 | 1.049 | 0.47 | 0.483 |
| Left preSMA RD | 3.336 | 66.644 | $< 10^{-15}$ | -0.169 | 0.866 | 1.792 | 1.717 | 0.866 |
| Right CST MD | 1.380 | 169.788 | $< 10^{-15}$ | 0.551 | 0.582 | 3.127 | 8.144 | $< 10^{-4}$ |
| Left CST MD | 2.289 | 85.986 | $< 10^{-15}$ | 0.564 | 0.573 | 2.738 | 4.446 | **0.004** |
| Right CST FA | 2.091 | 73.91 | $< 10^{-15}$ | -1.292 | 0.197 | 2.842 | 4.83 | **0.002** |
| Left CST FA | 2.739 | 55.692 | $< 10^{-15}$ | -1.156 | 0.248 | 2.803 | 2.603 | **0.03** |
| Right CST RD | 1.981 | 107.946 | $< 10^{-15}$ | 1.082 | 0.280 | 2.925 | 6.007 | **0.0006** |
| Left CST RD | 2.507 | 75.863 | $< 10^{-15}$ | 1.091 | 0.276 | 2.837 | 3.801 | **0.008** |
| Right caudate MD | 1.627 | 9.749 | **0.0002** | -2.489 | **0.013** | 3.117 | 6.339 | **0.0002** |
| Left caudate MD | 2.220 | 3.623 | **0.034** | -2.830 | **0.005** | 2.805 | 2.884 | **0.03** |
| Right putamen MD | 1.946 | 24.992 | $< 10^{-10}$ | 1.590 | 0.112 | 3.263 | 6.941 | $< 10^{-4}$ |
| Left putamen MD | 2.980 | 23.468 | $< 10^{-13}$ | 0.198 | 0.843 | 2.645 | 2.919 | **0.04** |

The SSRT<sub>test-retest</sub> variable was included in the models with SSRT variables. SSRT<sub>test-retest</sub> was significantly associated with SSRT, SSRT<sub>log</sub>, Go RT<sub>median</sub>, and Go RT<sub>STD</sub> ( $t < -7.632$ , $p < 10^{-13}$ ), but not with SSD50 ( $t = -1.514$ , $p = 0.131$ ). Abbreviations: SST: Stop-signal task, ROI: Region-of-interest, DTI: Diffusion tensor imaging, SSRT: Stop-signal reaction time, SSRT<sub>log</sub>: Logarithm-transformed SSRT, SSD 50: Stop-signal delay 50%. Go RT<sub>median</sub>: Median reaction time on correct go trials, Go RT<sub>STD</sub>: standard deviation for the go RT, IFG: inferior frontal gyrus; preSMA: pre-supplementary motor area; CST: corticospinal tract; DTI: diffusion tensor imaging; FA: fractional anisotropy, MD: mean diffusivity; RD: radial diffusivity, edf: estimated degrees of freedom.

Including handedness and average years of parent education as additional covariates in the above models did not significantly change the above age (F’s > 8.38, p’s < 10^-4^) or age-by-sex (F’s > 5.81, p’s < 0.0004) effects. Moreover, we did not observe any effects of parent education (t’s > −0.99, p’s > 0.32) or handedness (−1.07 < t’s < 0.50, p’s > 0.29) on any of the SST behavioral measures, except for the go RT_STD_, where higher average years of parent education was associated with less variable go RTs (t = −2.081, p = 0.038).

The developmental trajectories for SSRT for the initially good, intermediate and poor performers are plotted in Figure 5H-I. There was a significant difference in SSRT_log_ between groups (t = 8.393, p < 0.00001), as well as a significant age-by-group effect (t = −3.691, p = 0.0002), suggesting that the performance groups differed in their developmental trajectories of SSRT_log_. Initially good SSRT performers appeared to reach a developmental plateau around the age of 11 years, while the initially poor performers caught up with initially good performers around the age of 13-14 years (see Figure 5I). No age^2^-by-group effects were observed (t = 1.064, p = 0.288).

### 3.2. Age, sex and age-by-sex effects on the ROI microstructural measures

Age, sex and age-by-sex effects on the ROI DTI measures from the GAMM analyses are reported in Table 2. Spaghetti plots of the ROI DTI measures overlaid with GAMM estimated age trajectories for males and females are shown in Figure 6. Significant age-related changes were found for all ROIs, with FA increasing with age, and RD and MD decreasing with age. We did not observe any significant effects of sex on any of the ROI DTI measures, except for the left and right caudate nuclei, where females had lower MD than males. Moreover, significant age-by-sex interaction effects were observed for all ROI DTI measures, except for left IFG_FA_ and IFG_RD_, left preSMA_FA_ and preSMA_RD_, and right preSMA_FA_. Generally, males displayed more linearly increasing or decreasing maturational trajectories, while females generally displayed more curvilinear maturational trajectories that increased or decreased more steeply in the early years of the included age range and started to level off around the age of 12 to 14 years.

**Figure 6.**
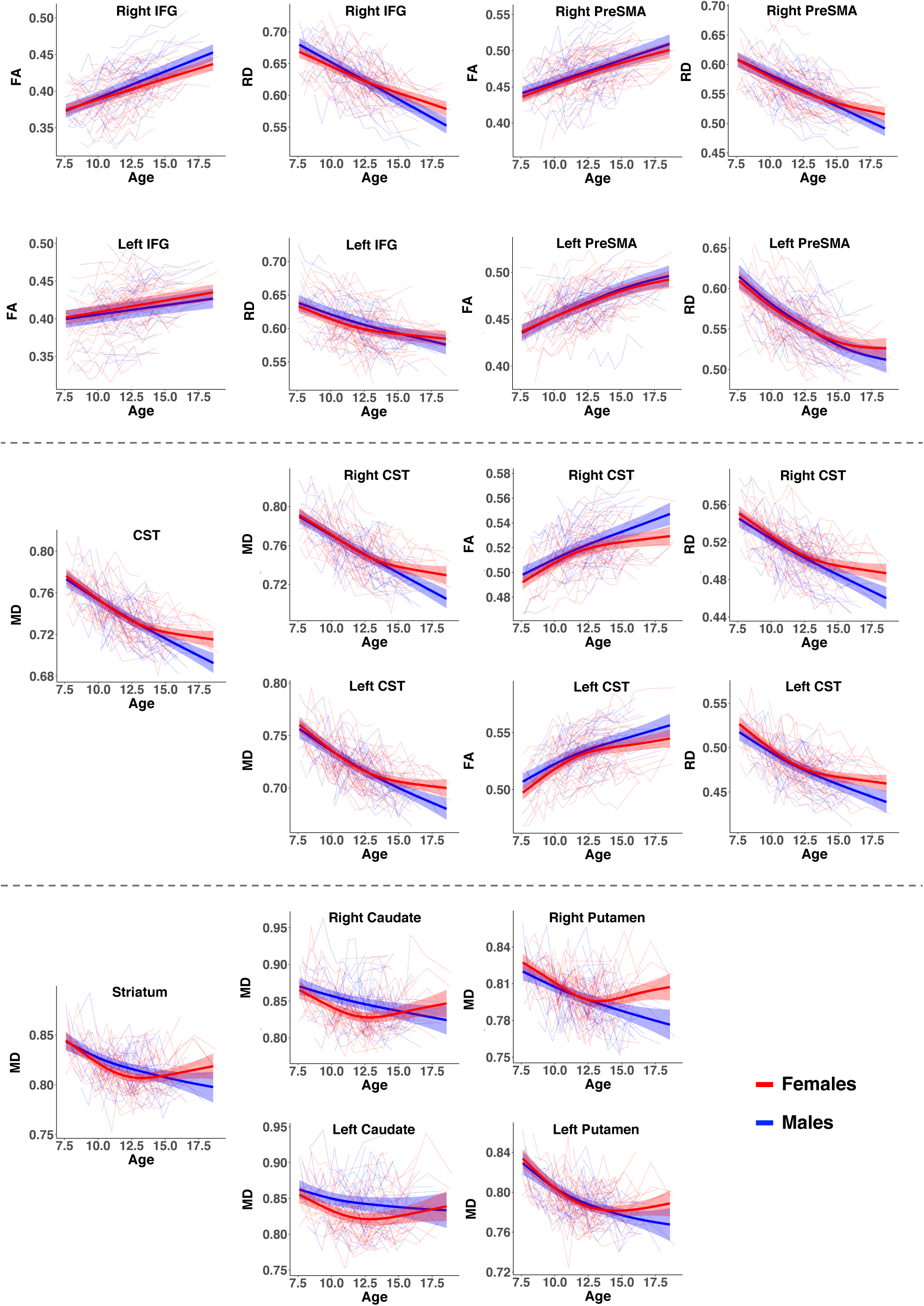
Spaghetti plots of ROI DTI measures overlaid with GAMM estimated age trajectories with shaded 95% confidence intervals for males (blue) and females (red). From top to bottom: left and right inferior frontal gyrus (IFG) fractional anisotropy (FA) and radial diffusivity (RD), left and right pre-supplementary motor area (preSMA) FA and RD, bilateral corticospinal tracts (CST) mean diffusivity (MD) and left and right CST MD, FA and RD, neostriatum MD and left and right caudate and putamen MD. Values for MD and RD are given in 10^−3^ mm^2^/s.

Including movement during DWI, average years of parent education and handedness as additional covariates in the above models, did not significantly change the age (F’s > 7.969, p’s < 0.0003) or age-by-sex effects (F’s > 2.69, p’s < 0.03), reported above. The lack of an effect of sex remained, except for right putamen MD (t = 2.141, p = 0.033) where females had higher MD than males, and left caudate MD (t = −2.515, p = 0.012), where females had lower MD than males. The previous observed sex effect for right caudate MD became non-significant (t = −1.944, p = 0.052). More movement during DWI was significantly associated with higher FA and lower RD and MD in all ROIs (−3.576 > t’s > 3.326, p’s < 0.0004), except for right preSMA_FA_ and right and left CST FA (−0.808 < t’s < 0.907, p’s > 0.07). Additionally, movement during DWI decreased significantly and linearly with age (edf = 1.026, F = 80.202, p < 10^-15^), but was not significantly associated with sex (t = −0.767, p = 0.443) or age-by-sex (edf = 1.003, F = 2.438, p = 0.119). Moreover, higher average years of parent education was related to higher right preSMA_FA_, lower RD in right IFG and preSMA, and left and right CST, and lower MD in bilateral CST, striatum, left and right CST, and right putamen (−1.978 > t’s > 2.980, p’s < 0.05). We did not observe any effects of handedness, except for right putamen MD (t = −1982, p = 0.048), where right-handed participants had higher MD than left-handed.

### 3.3. Associations between SSRT_log_ and right IFG and right preSMA FA and RD

Results concerning our main hypotheses that developmental improvements in SSRT would be associated with individual differences in the maturational trajectories of FA and RD of the white matter underlying the right IFG and right preSMA, are reported in Table 3. We did not observe any significant associations between SSRT_log_ and the age-by-right IFG_FA_, -right IFG_RD_ or -right preSMA_RD_ interactions (Table 3, models 1, 3 and 4), although there was some indication for a possible age-by-right preSMA_RD_ effect (p = 0.035). Moreover, there were no significant main effects of ROI microstructure. We did, however, find a significant age-by-right preSMA_FA_ effect that survived correction for multiple comparisons (Table 3, model 2). Children with lower right preSMA_FA_ exhibited poorer SSRT performance at younger ages and steeper developmental trajectories of SSRT_log_ relative to children with higher right preSMA_FA_ (Figure 7A, C and D). This interaction effect remained significant (β = 2.991, t = 2.799, p = 0.005), when additionally controlling for movement during DWI, average years of parent education and handedness. Moreover, the age-by-right preSMA_FA_ interaction effect remained significant (β’s > 2.951, t’s > 2.748, p’s < 0.006), when adding whole skeleton FA or left preSMA_FA_ and their interactions with age to the latter model, suggesting that our finding is anatomically specific. Of note, splitting right preSMA_FA_ into initial high, medium and low FA groups, using the same approach for FA as used for SSRT in section 2.10.3, showed that the three FA groups differed in average FA levels (main effect of group: t = 15.628, p < 0.0001) but exhibited similar maturational trajectories (Figure 7B), i.e. no significant age- or age^2^-by-group interaction effects (p > 0.590).

**Figure 7.**
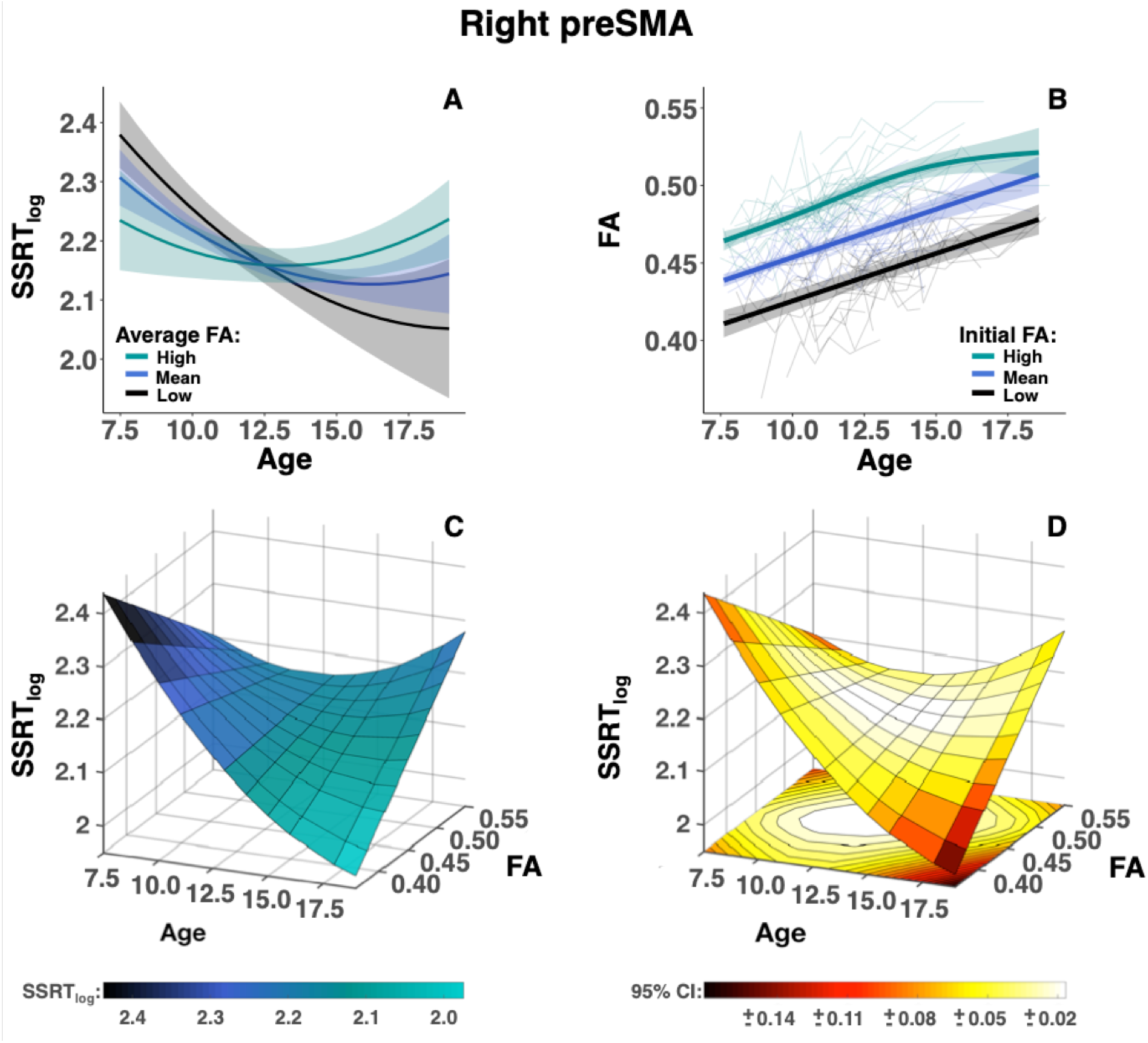
Depiction of the observed significant age-by-right preSMA FA effect on SSRT_log_ in a 2D (A) and 3D (C) plot based on the fit from the hypothesis model. In (A), the fitted developmental trajectories of SSRT_log_ with shaded 95% confidence intervals are depicted for high (+2 standard deviation, cyan), mean (blue) and low (xy12 standard deviation, black) average right preSMA FA. (B) depicts a spaghetti plot of right preSMA FA overlaid with GAMM estimated age trajectories with shaded 95% confidence intervals for individuals with initial low (black), medium (blue) and high (cyan) right preSMA FA. The three FA groups differ significantly in average FA levels but exhibit similar maturational trajectories. (C) depicts the association between SSRT_log_, age and right preSMA FA in 3D with the surface color-coded by the SSRT_log_ values. (D) represents the 95% confidence intervals (CI) that can be derived by adding or subtracting the values represented in the color bar from the SSRT_log_ values at a given square in the 3D sheet.

**Table 3.** Results from the a priori models predicting SSRT_log_

| Model | SST variable | ROI |  |  | ROI-by-age |  |  |
| --- | --- | --- | --- | --- | --- | --- | --- |
| | | $\beta$ | t | p | $\beta$ | t | P |
| 1 | Right IFG FA | 0.029 | 0.486 | 0.627 | 1.266 | 1.029 | 0.304 |
| 2 | Right preSMA FA | -0.029 | -0.551 | 0.582 | 3.186 | 2.972 | **0.003** |
| 3 | Right IFG RD | -0.085 | -1.462 | 0.144 | -1.681 | -1.349 | 0.178 |
| 4 | Right preSMA RD | -0.027 | -0.487 | 0.626 | -2.567 | -2.114 | 0.035 |

All models included $SSRT_{test-retest}$ , age and age<sup>2</sup>. Furthermore, model 1 also included sex and age-by-sex, while models 3 and 4, additionally included sex, age-by-sex, and age<sup>2</sup>-by-sex (data not shown). In all models, $SSRT_{log}$ was significantly associated with $SSRT_{test-retest}$ , age and age<sup>2</sup> ( $p$ 's < 0.017). In Model 1, 3 and 4, there were no significant effects of sex or age-by-sex or age<sup>2</sup>-by-sex (models 3 and 4 only) ( $p$ 's > 0.361). Abbreviations: $SSRT_{log}$ : logarithm transformed stop-signal reaction time; SST: stop signal task; IFG: Inferior frontal gyrus, PreSMA: Pre-supplementary motor area; FA: Fractional anisotropy, RD: Radial diffusivity

To further investigate the nature of the age-by-right preSMA_FA_ interaction, we also examined for a possible age-by-right PreSMA_AD_ interaction effect on SSRT_log_. Based on a GAMM model investigating age, sex and age-by-sex effects on right PreSMA_AD_ (age: edf = 1.014, F = 88.987, p < 10^-15^; sex: t = −0.765, p = 0.444; age-by-sex: edf = 2.449, F = 5.332, p = 0.003), we tested the following LME model: SSRT_log_ = SST_test-retest +_ age + age^2^ + sex + age*sex + age^2^*sex + ROI + age*ROI. We did not observe any significant effects for right PreSMA_AD_ (β = −0.053, t = −1.048, p = 0.295) or age-by-right PreSMA_AD_ (β = 0.998, t = 0.886, p = 0.376). Overall, our findings suggest that the observed age-by-right preSMA_FA_ interaction effect on SSRT_log_ was mainly driven by RD and less so by AD, with children with higher right preSMA_RD_ and lower AD exhibiting steeper developmental trajectories of SSRT_log_ relative to children with lower right preSMA_RD_ and AD.

Finally, exploratory analyses of left IFG_FA_ (model: SSRT_log_ = SST_test-retest +_ age + age^2^ + ROI + age*ROI), and left IFG_RD_ and left preSMA_FA_ and preSMA_RD_ (model: SSRT_log_ = SST_test-retest +_ age + age^2^ + ROI + age*ROI + age^2^*ROI) did not reveal any significant main effects of ROI microstructure (p’s > 0.750) or ROI-by-age or age^2^ interaction effects (p’s > 0.120) on SSRT_log_.

### 3.4. Associations between SSRT_log_ and CST and striatum DTI measures

Exploratory analyses concerning CST MD did not reveal any significant main effect of bilateral CST MD or age- or age^2^ -by-bilateral CST MD interaction effects on SSRT_log_ (p’s > 0.208), nor right CST MD, FA, or RD (p’s > 0.172). There was some indication of an association between SSRT_log_ and left CST MD (t = 1.861, p = 0.063), FA (t = −1.819, p = 0.070) and RD (t = 2.023, p = 0.044). However, when adding movement during DWI, handedness and average years of parent education in the models, these associations disappeared for left CST MD and RD (MD: t = 0.637, p = 0.524, RD: t = 1.248, p = 0.213), but not FA (t = −1.735, p = 0.083). Exploratory analyses of bilateral striatum did not reveal any significant associations with SSRT_log_ for striatum MD or age- or age^2^-by-striatum MD (p’s > 0.211). Likewise, no significant associations were observed between SSRT_log_ and left or right caudate nucleus or putamen or age- or age^2^-by-left or right caudate nucleus or putamen MD interactions (p’s > 0.214). Finally, we did not observe any significant sex, age-by-sex or age^2^-by-sex effects on SSRT_log_ for bilateral CST MD, left and right CST MD, FA or RD, or striatal or left and right caudate or putamen MD (p’s > 0.258).

### 3.5. Associations between SSRT_log_ and age-by-FA across the white matter skeleton

An effect size map of the association between SSRT_log_ and age-by-FA across the white matter skeleton is presented in Figure 8. The full t-map can be downloaded from https://neurovault.org/collections/AMYMNIEX/. Positive associations (t-value ≥ 1.965, p ≤ 0.05, uncorrected) were observed in several clusters across the white matter skeleton, including the left forceps minor (slice 52), the right anterior corona radiata/inferior fronto-occipital fasciculus (slice 31), the white matter underlying the left (slice 15) and right preSMA (slice 9) and the right motor cortex (slices −1 and −7), the left posterior limb of the internal capsule (slices −1, −7 and −12), the right CST (slice −22), the left superior longitudinal fasciculus (slice −42), the bilateral white matter laterally to the forceps major; a region containing a mixture of different tracts, such as the posterior thalamic radiation and the optic tract (slice −57), as well as the right side of the forceps major (slice −79). Overall, SSRT_log_ was positively and negatively associated with age-by-FA in respectively, 6.8% and 0.4% of all skeleton voxels (1.965 ≤ t ≤ −1.965, p ≤ 0.05, uncorrected). In comparison, SSRT_log_ was positively associated with age-by-FA in 19.5% of right preSMA ROI voxels, 6.4% of the left preSMA ROI voxels, 2.6% and 0% of respectively the right and left IFG ROI voxels, and 4.2% and 6.8% of respectively the right and left CST ROI voxels. None of the voxels within the left and right preSMA and right IFG ROIs showed negative associations with SSRT_log_. However, negative associations were observed in 2.7%, 0.6% and 0.4% of the voxels of respectively the left IFG, right CST and left CST ROIs.

**Figure 8.**
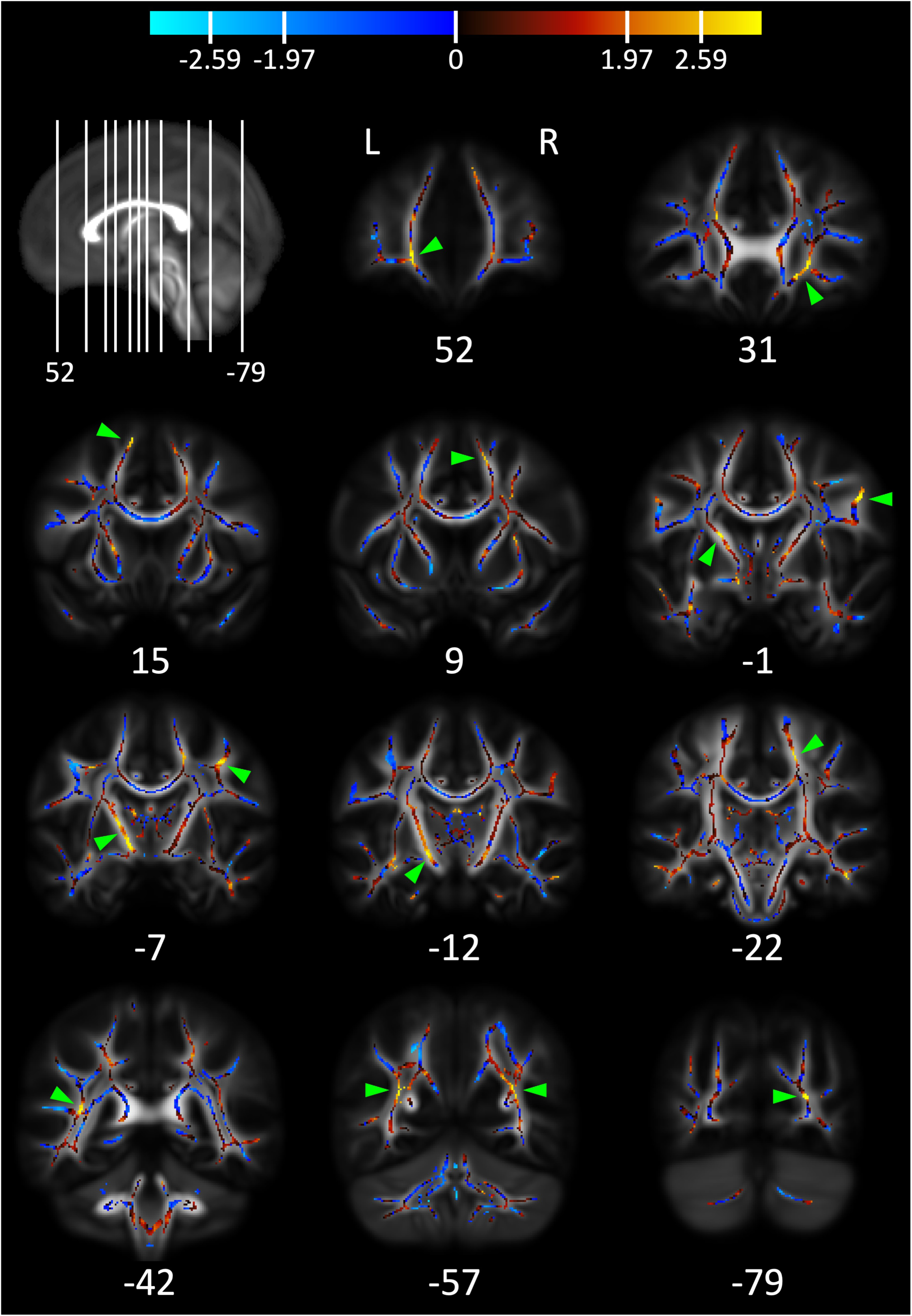
Effect size map displaying the association between SSRT_log_ and age-by-FA across the white matter skeleton, after correction for SST_test-retest_, movement during DWI, age, age^2^, sex, age-by-sex, age^2^-by-sex and FA. The map displays the images with the highest t-values and largest clusters, based on visual inspection of the effect size map, with the main clusters of interest pointed out using green arrows (see section 3.5. for more details). The color bar shows the color mapping of the skeleton voxels to the values of uncorrected t-statistics. The positive and negative t-values in the color bar correspond to t = ±1.97, p = 0.05 and t = ±2.59, p = 0.01 (df = 480, two-tailed, uncorrected). The MNI Y coordinates for the coronal slices are given under each image. In the top left corner, the depicted coronal slices are shown on a midsagittal image, in which the most anterior line represent slice 52 and the most posterior line represent slice −79. Images are shown according to neurological convention, where L = left hemisphere, R = right hemisphere.

## 4. Discussion

In the present longitudinal study in typically-developing children and adolescents aged seven to 19 years, assessed up to nine times, we investigated whether individual differences in developmental improvements in motor response cancellation were associated with the maturational trajectories of the white matter microstructure underlying the right IFG and the right preSMA. Motor response cancellation performance, estimated by SSRT, improved non-linearly with age, and appeared to reach a developmental plateau around the age of 13-14 years. Moreover, initially good and poor performers exhibited different developmental trajectories of SSRT improvement. As hypothesized, we found a preSMA_FA_ FA-by-age interaction effect on SSRT, where children with lower right preSMA_FA_ exhibited steeper developmental trajectories of SSRT improvement relative to children with higher right preSMA_FA_. This relationship appeared to be anatomically specific and not driven by individual differences in global white matter FA. Further, the relationship did not appear to be mediated by individual differences in parent education, handedness or subject movement during DWI scanning. We did not observe any statistically significant relationships between improvements in SSRT and the maturational trajectories of right IFG FA or RD or right preSMA_RD_, nor did we find any associations with CST or striatum MD in exploratory analyses.

### 4.1. Development of SSRT performance

All SST measures improved with age. The significant improvement with age in cancelling motor responses, i.e. faster SSRT, appeared to reach a plateau around the age of 13-14 years. Our observation of faster SSRT with increasing age during childhood generally agrees with existing literature (Curley et al., 2018; Dupuis et al., 2019; Williams et al., 1999). Findings from two cross-sectional studies (Williams et al., 1999), though, suggest that age-related improvements in SSRT may continue a few more years into adolescence than observed in the present study. However, in a recent longitudinal study using the same stop-signal task as in our study, the improvement in SSRT seemed to level off toward the end of the investigated age range of four to 13 years (Curley et al., 2018), suggesting similar developmental trajectories of SSRT as observed in our study. These discrepancies in when SSRT reaches a developmental plateau may be due to differences in e.g. task design, cohort characteristics, or the use of cross-sectional vs. longitudinal study designs. For example, the test-retest effects observed in the current study may have accelerated the magnitude of improvement above and beyond “pure” age-related improvements, thereby potentially causing participants to reach their developmental plateau at an earlier age. The latter would be independent of our correction for test-retest effects that ensured that the magnitude of improvement between the first and second assessment was not overestimated (Hoffman et al., 2011; Sullivan et al., 2017). Interestingly, we observed that initially good, intermediate and poor performers significantly differed in their developmental trajectories of SSRT performance. Initially good performers reached a plateau around the age of 11 years, while initially poor performers showed a more protracted developmental trajectory that plateaued around the age of 14 years. Moreover, while initially good and poor performers differed in SSRT performance at younger ages, poor performers displayed performance levels comparable with good performers around the age of 13-14 years. Combined, these findings suggest that initially poor performers’ SSRT development may be delayed, potentially due to individual differences in the rate or phase of SSRT development, or experience-dependent plasticity. These results highlight the importance of longitudinal observations for differentiating between possible stable and dynamic patterns of change.

### 4.2. Maturational trajectories of ROI microstructure

Our findings that in all ROIs FA increased and RD and MD decreased with age are in agreement with the existing literature (Lebel et al., 2019). The validity of the observed sex differences is less clear. Females had lower left and right caudate MD than males. Furthermore, while males generally displayed more linear maturational trajectories, females had more curvilinear maturational trajectories that changed more steeply in the early years of the included age range and leveled off around the age of 12-14 years. As reviewed previously (Lebel et al., 2019; Tamnes et al., 2018), there are a few studies that also observed more prolonged maturation of diffusion measures in males as compared to females. While it has been hypothesized that such a pattern may be related to earlier puberty in females, available evidence is inconclusive (Lebel et al., 2019). In general, only few studies have examined sex differences in white matter and subcortical grey matter microstructure during childhood and adolescence, and findings have generally been inconsistent (Kaczkurkin et al., 2019; Tamnes et al., 2018). Moreover, most studies examining white matter microstructural maturation have been conducted in cross-sectional samples (Lebel et al., 2019). Of nine longitudinal studies, i.e. (Herting et al., 2017; Oyefiade et al., 2018) and the seven reviewed in (Lebel et al., 2019), only Simmonds et. al. (2014) acquired more than two scans on average (mean = 2.5), with approximately half of the typically-developing participants aged eight to 29 years having been scanned three to five times. In comparison, in the current study participants were assessed on average 6.6 times, with only five of the 88 participants having only two timepoints. This high number of timepoints, as well as use of a robust non-parametric smooth function to estimate age trajectories (Fjell et al., 2010), should, at least in principle, yield an improved model of underlying maturational changes.

### 4.3. Associations between developmental SSRT improvement and preSMA microstructure

We observed that children and adolescents with lower average right preSMA_FA_ across the individual maturational trajectory displayed lower motor cancellation performance at younger ages and exhibited steeper developmental trajectories of SSRT improvement. Children and adolescents with higher average right preSMA_FA_ exhibited flatter developmental trajectories of SSRT improvement along with faster SSRT already at the first assessments. This effect was mainly driven by higher right preSMA_RD_ for children with lower right preSMA_FA_ and lower right preSMA_RD_ for children with higher right preSMA_FA_. In developing children and adolescents, this pattern of increased FA and decreased RD is typically associated with a more mature brain (Lebel et al., 2019). While the interpretation of DTI measures is not straight forward, increased FA and decreased RD have previously been linked to e.g. increased axon density, diameter, or myelination as well as decreased levels of crossing fibers (Beaulieu, 2009; Schwartz et al., 2005). Furthermore, we found that children with initially high, medium and low right preSMA_FA_ exhibited similar linear maturational trajectories in FA but differed in average FA levels. This suggests that, among typically-developing children, there may be stable individual differences in the underlying neural architecture that persist despite ongoing maturation of the white matter, at least within the investigated age range. Intriguingly, while right preSMA_FA_ continued to increase linearly throughout the investigated age range, SSRT performance levelled off around the age of 13-14 years. It is unclear what mechanisms may underly the observed brain-behavioral developmental relationship pattern and the effects may not be causal. It may be that when the right preSMA white matter reaches a certain level of maturity, as reflected in tissue FA, it is sufficient to support maximum SSRT performance levels and, thus, no further improvement in motor response cancellation is achieved. Furthermore, since SSRT performance involves an extended brain network, it may also be that other parts of the network (Aron, 2011) contribute to the observed pattern.

### 4.4. SSRT and right IFG microstructure

Contrary to our expectations, we did not observe significant associations between developmental improvements in SSRT and maturational changes in FA or RD of the white matter underlying the right IFG, nor did we find any main effects of right IFG FA or RD (i.e. independent of age). This observation is in apparent contrast to our baseline study of the same cohort, where faster SSRT was associated with higher FA and lower RD in the white matter underlying the right IFG (Madsen et al., 2010). Moreover, a recent longitudinal study in typically-developing children aged four to 13 years using the same stop-signal task found that faster SSRT was associated with relatively larger cortical surface area of the bilateral, and especially the right, opercular region of the IFG (Curley et al., 2018). Notable, the latter study did not observe any associations between age-related changes in SSRT performance and changes in IFG cortical surface area, suggesting that the association between SSRT and IFG cortical surface area were mediated by stable differences in IFG cortical surface area between individuals. Cortical surface area generally peaks in pre-adolescence (Bogg et al., 2012), while DWI measures generally continue to mature into adolescence and early adulthood (Lebel et al., 2019). Furthermore, the relationship between cortical surface area and white mater microstructure is not clear. Nevertheless, our finding seems to suggest that maturational changes in the white matter underlying right IFG may not play a large role in improved motor response cancellation performance with age. In light of the lack of similar studies, our failure to observe effects for the right IFG should be considered preliminary and needs confirmation in future longitudinal studies.

### 4.5. SSRT and CST and neostriatal microstructure

Our exploratory analyses of possible associations between developmental improvements in SSRT and maturational changes in CST, putamen and caudate microstructural measures did not reveal any significant associations. However, we cannot draw any strong conclusions from these findings. It might be that our measures were anatomically too nonspecific (Guo et al., 2018), as we did not, for example, differentiate between motor, cognitive or emotional areas in the caudate nucleus or putamen (Draganski et al., 2008). Also, the skeleton segments included in the CST ROIs extended towards both the motor and sensory motor cortices. Interestingly, the effect size maps depicting the age-by-FA interaction effects on the developmental changes in SSRT performance across the white matter skeleton revealed a positive association in the posterior limb of the internal capsule, suggesting that this part of the CST might play a role in developing SSRT performance.

### 4.6 Possible limitations

We investigated a limited age range between seven and 19 years with the ages at the extremes being underrepresented. Furthermore, there were relatively fewer males than females. Thus, we cannot infer what happens at younger or older ages or generalize observed sex differences or the lack thereof. Moreover, even though we employed an adaptive stop-signal task, where task difficulty is dynamically adjusted to minimize ceiling effects, we cannot rule out a possible link between task difficulty and observed plateaus in SSRT performance. Furthermore, while we used state of the art image processing pipelines and corrected for e.g. movement during DWI scanning, we cannot rule out that individual processing steps, or combinations of these, may have biased our results. Nevertheless, given the increased statistical power and sensitivity provided by the high number of time points per subject and our use of robust non-parametric smooth functions, we believe that the observed developmental changes in SSRT performance and the maturational trajectories in ROI microstructure as well as their interrelations contribute valuable additional information about maturing motor inhibitory functions. Finally, due to the observational nature of our study, we cannot infer causality.

## 5. Conclusions

Children displayed different developmental trajectories of motor response cancellation performance. Initially well performing children plateaued around the age of 11 years, while initially poor performers first caught up three years later. As hypothesized, developmental improvements in motor response cancellation performance were reflected in the maturational trajectories of the white matter microstructure underlying the right preSMA. Children with lower average right preSMA_FA_ exhibited poorer SSRT performance at younger ages and steeper developmental trajectories of SSRT improvement. Children with higher average right preSMA_FA_ exhibited flatter developmental trajectories of SSRT. While SSRT performance generally levelled off around the age of 13-14 years, right preSMA_FA_ continued to increase linearly throughout the investigated age range, with apparent stable differences in FA levels between children with initial low, medium or high right preSMA_FA_. It is unclear what may underlie this pattern of brain-behavior developmental association. It may be that once a certain level of maturity in the white matter underlying the right preSMA is reached, it is sufficient to support maximum SSRT performance levels and no further improvement in motor response cancellation is achieved. Similar dynamics may apply to other behavioral read-outs and brain structures and, thus, need to be considered in longitudinal MRI studies that are designed to map brain structural correlates of behavioral changes during development.

## Acknowledgements

The authors sincerely thank the children and their parents for their participation in the HUBU study.

